# Siderophore piracy enables the nasal commensal *Staphylococcus lugdunensis* to antagonize the pathogen *Staphylococcus aureus*

**DOI:** 10.1101/2024.02.29.582731

**Authors:** Ralf Rosenstein, Benjamin O. Torres Salazar, Claudia Sauer, Simon Heilbronner, Bernhard Krismer, Andreas Peschel

## Abstract

Bacterial pathogens such as *Staphylococcus aureus* colonize body surfaces of part of the human population, which represents a critical risk factor for skin disorders and invasive infections. However, such pathogens do not belong to the human core microbiomes – beneficial ‘commensal’ bacteria can often prevent the persistence of certain pathogens – using molecular strategies that are only superficially understood. We recently reported that the commensal bacterium *Staphylococcus lugdunensis* produces the novel antibiotic lugdunin, which eradicates *S. aureus* from nasal microbiomes of hospitalized patients. However, it has remained unclear if *S. lugdunensis* may affect *S. aureus* carriage in the general population and how *S. lugdunensis* carriage could be promoted to enhance its *S. aureus*-eliminating capacity. We could cultivate *S. lugdunensis* from the noses of 6.3% of healthy human volunteers. In addition, *S. lugdunensis* DNA could be identified in metagenomes of many culture-negative nasal samples indicating that cultivation success depends on a specific bacterial threshold density. Healthy *S. lugdunensis* carriers had a 5.2-fold lower propensity to be colonized by *S. aureus* indicating that lugdunin can eliminate *S. aureus* also in healthy humans. *S. lugdunensis*-positive microbiomes were dominated by either *Staphylococcus epidermidis*, *Corynebacterium* species, or *Dolosigranulum pigrum*. These and further bacterial commensals, whose abundance was positively associated with *S. lugdunensis*, promoted *S. lugdunensis* growth in co-culture. Such mutualistic interactions depended on the production of iron-scavenging siderophores by supporting commensals, and on the capacity of *S. lugdunensis* to import siderophores. These findings underscore the importance of microbiome homeostasis for eliminating pathogen colonization. Elucidating mechanisms that drive microbiome interactions will become crucial for microbiome-precision editing approaches.

## Background

Host-associated microbiomes are shaped by mutualistic or antagonistic interactions among microbiome members. Some of these members depend on each other because they collaborate in the degradation and utilization of complex nutrient sources or the exchange of essential cofactors [1]. These processes can be of mutual benefit for two partners or can support only one of them, which may exploit other bacterium’s metabolic capacities. Interacting microorganisms can also antagonize each other directly by the release of antimicrobial bacteriocins, which inhibit major competitors to enhance the producer’s fitness [2]. Such mutualistic and antagonistic mechanisms govern complex, often multi-dimensional interaction networks, which have remained largely unexplored.

Elucidating mechanisms that can promote the persistence of beneficial and impair that of harmful bacteria in microbiomes is of particular relevance for the prevention of infections caused by bacterial pathogens that use human microbiomes as their major reservoirs [3]. All of the notoriously antibiotic-resistant pathogens, vancomycin-resistant *Enterococcus faecalis* and *Enterococcus faecium*, methicillin-resistant *Staphylococcus aureus* (MRSA), and carbapenemase- or extended-spectrum beta-lactamase-producing *Klebsiella pneumoniae*, *Acinetobacter baumannii*, and *Escherichia coli*, can be found in the microbiomes of healthy humans or of at-risk patients [4]. Carriage of such facultative antibiotic-resistant pathogens strongly increases the risk of invasive, difficult-to-treat infections [5]. Microbiome composition has a dominant role in the capacity of pathogens to colonize, most probably as a consequence of antagonistic effects exerted by beneficial commensals [6]. However, current options for decolonization of pathogens are very limited, which demands the development of effective and specific pathogen eradication regimes, which should maintain microbiome integrity [7].

*S. aureus* colonizes the nares of 30% to 40% of the human population, which represents a major risk factor for severe *S. aureus* infections, in particular when caused by MRSA strains [8, 9]. The abundance of IgA in the nose has been found to shape bacterial densities in the nose but not the presence or absence of *S. aureus* or other nasal species [10]. The nasal microbiome is highly diverse, and its composition is crucial for the capacity of *S. aureus* to colonize [11]. Nevertheless, we are only beginning to understand the mechanisms used by nasal commensals to exclude *S. aureus*. Human nasal microbiome analyses have defined at least seven different community state types (CSTs), which are dominated by specific bacterial signature species or groups [10, 11].

We recently reported that the production of bacteriocins is very frequent among commensal bacterial species of the human nasal microbiomes suggesting that such mechanisms should have a strong impact on microbiome composition and, potentially, the capacity to exclude specific pathogens [12, 13]. Notably, many of these compounds did not or not only act against closely related species, which contradicts previous definitions of bacteriocins’ roles and indicates that bacteriocins are often produced to combat major competitors, irrespective of the relatedness of bacteriocin producer and target strain [2]. One of the antimicrobial molecules, the novel fibupeptide lugdunin, produced by the coagulase-negative *Staphylococcus* (CoNS) species *Staphylococcus lugdunensis*, was explored in detail [14]. Almost all of the nasal *S. lugdunensis* isolates contained the lugdunin-biosynthetic gene cluster in the chromosome. *S. lugdunensis* eradicated *S. aureus* in a lugdunin-dependent fashion, during cultivation in laboratory media and in animal models. Moreover, hospitalized patients carrying *S. lugdunensis* in their noses had a six-fold lower risk to be colonized by *S. aureus*. However, only 9.1% of the hospitalized patients were *S. lugdunensis* carriers, raising the question why only certain nasal microbiomes may permit the persistence of *S. lugdunensis* with its *S. aureus*-eradicating activity [14]. Here we describe that *S. lugdunensis* is equally rare in healthy humans as in hospitalized patients and promotes the elimination of *S. aureus* in both groups of humans. *S. lugdunensis* was positively associated with several other commensals, which were required for *S. lugdunensis* to thrive by providing essential iron-scavenging siderophores.

## Methods

### Bacterial strains and media

Bacterial strains used in this study are summarized in Suppl. Table 3. All bacteria were grown on basic medium (BM; 1% soy peptone, 0.5% yeast extract, 0.5% NaCl, 0.1% glucose and 0.1% K_2_HPO_4_, pH 7.2), which was supplemented with 5% sheep blood (Oxoid) and 1.5% agar (BD European Agar) if needed. *Cutibacterium acnes* and *Lawsonella clevelandensis* were incubated under anerobic conditions using an anaerobic jar and AnaeroGen^TM^ (Thermo). For the phenotypic identification of *S. lugdunensis*, *S. aureus* and other staphylococcal species in nasal swabs, basic medium (BM), blood agar and selective agar SSL [15] were used. For the siderophore experiments, bacteria were grown in E-BHI, composed of brain heart infusion medium (BHI; Roth) supplemented with 10 µM of the iron chelator ethylenediamine-di(o-hydroxyphenylacetic acid) (EDDHA; LGC Standards GmbH), for 72 h at 37°C and constant shaking at 160 rpm. For transformation experiments with *E. coli* DC10B and *S. lugdunensis*, tryptic soy broth (TSB; Oxoid) or tryptic soy agar (TSA) were used and, when necessary, supplemented with antibiotics at concentrations of 10 µg/ml chloramphenicol (Sigma-Aldrich), 100 µg/ml ampicillin (Roth) and 1 µg/ml anhydrotetracycline (Fluka).

### Human volunteer selection, nasal swabbing, and detection of *S. lugdunensis* and *S. aureus*

The sample collection procedures were approved by the clinical ethics committee of the University of Tübingen (No. 109/2009 BO2) and informed written consent was obtained from all volunteers. Nasal swabs were taken exclusively from healthy adults. Samples from 270 healthy volunteers of the University of Tübingen were collected by swabbing both nares with cotton swabs and suspending them in 1 ml phosphate buffered saline (PBS). Various dilutions of each sample were plated on BM, blood agar, and selective *S. lugdunensis* medium agar (SSL) [15] for phenotypic identification of *S. lugdunensis*, *S. aureus* and other staphylococcal species. The plates were incubated for 24 to 48 h at 37°C under aerobic and anaerobic conditions, respectively. The bacterial identity was evaluated by matrix-assisted laser desorption/ionization-time-of-flight mass spectrometry (mass spectrometer: AXIMA Assurance, Shimadzu Europa GmbH, Duisburg, database: SARAMIS with 23.980 spectra and 3.380 superspectra, BioMérieux, Nürtingen).

### Metagenome sequencing of nasal microbiome samples

Nasal microbiome samples were taken from individuals by swabbing both nares successively with one nylon-flocked E-Swab (ThermoFisher Scientific) and suspending them in 1 ml Amies transport medium. Two replicate swabs were consecutively taken per volunteer. For degradation of contaminating host DNA and subsequent preparation of bacterial DNA, the suspended samples were mixed with AHL-buffer and treated with the QIAamp DNA Microbiome Kit (Qiagen) according to the manufacturer’s instructions. Finally, DNA was eluted from the spin columns in 50 µl AVE-buffer. DNA concentration was determined with Qubit 3.0 (Thermo Fisher Scientific) using High Sensitivity (HS) reagents for low DNA concentrations. 2.5 µl of the preparation was used for whole-genome amplification with REPLI-g Single Cell Kit (Qiagen). Subsequently, the amplified DNA was purified with Genomic DNA Clean & Concentrator (Zymo Research) and the final DNA concentration was determined with Qubit 3.0.

About 10 µg of the purified DNA was used for next generation sequencing by GATC (Konstanz, Germany). In brief, libraries of fragments of ca. 400 bases were prepared and submitted to Illumina paired-end sequencing with read lengths of 150 bases. After quality-filtering, an average of 30 million read pairs was obtained per sample and provided as compressed fastq.gz file (separate files for each end of the sequenced inserts).

In the first metagenome analysis (after 18 months) we obtained a median of 70.13 million quality-filtered, paired-end reads per sample. In the second metagenome analysis (after 23 months), a median of 52.71 million reads per sample were obtained. Despite taking measures to reduce the proportion of human DNA in the samples we found varying percentages of metagenomic reads mapping to the human DNA reference database: at the first time point (18 months), human-derived reads ranged from 14.4% to 94.3% (median: 74.75%) per sample while at time point 2 (23 months) between 0.5% and 90.9% (median: 28.15%) of the reads mapped to sequences from the hg19 database. At time point 1, a median of 11.45 million bacterial reads per sample (sample D had exceptionally low read numbers due to antibiotic treatment and was left out as an outlier) were obtained, while at the second time point the median of bacterial reads was 21.81 million.

### Bioinformatics

The sequence reads from each file were mapped against the NCBI non-redundant (nr) protein database by using the optimized BLASTX algorithm of DIAMOND [16]. For the subsequent functional and taxonomic analysis with MEGAN6 [17], the DIAMOND alignment files were “meganized” by combining the paired-end sequences for each sample and providing them with functional annotations. Taxonomic binning, functional analysis, and statistical analyses were performed by using the corresponding functions of MEGAN6. For the comparison of the metagenomes on the genus level, pairwise distances were calculated based on ecological distance according to Hellinger [18].

To exploit the genetic information obtained with the metagenome sequence reads we assembled the species-specific reads into larger contiguous sequences. To this end, we used the MetaSpades tool, which is implemented in the SPADES assembler [19]. For species-specific read assemblies, we first used MEGAN6 to extract the reads assigned to the species of interest as a Fasta file. Then we used MetaSpades to assemble these reads into contigs.

In order to determine the *S. lugdunensis* strains that reside in the anterior nares of the carriers we selectively extracted the *S. lugdunensis* reads where possible and assembled them into contigs. The contigs were compared with a database composed of *S. lugdunensis* genome sequences by BlastN and the genomes getting the highest number of hits were regarded as the closest relative(s) to the strains present in the *S. lugdunensis* carriers. Thus, we obtained strain profiles based on the total alignment lengths of hits mapped to a phylogenetic tree of *S. lugdunensis* strains (Fig. 2).

Correlation assays were performed by network analysis with the web-based tool MetagenoNets, (https://web.rniapps.net/metanets/; [20]). To this end, an abundance table with read numbers of the bacterial species in the samples was uploaded to MetagenoNets. The abundance data were transformed to centered log-ratios, filtered for a prevalence (minimum proportion of reads in samples) of 0.1%, and an occurrence (minimum percentage of samples showing the species) of 20%. Network inference was calculated by NAMAP (a modified ReBoot method; [21, 22]) based on Pearson correlation coefficients. The results were downloaded as correlation matrix, imported into R [23] using the package corrr [24] and visualized with the R package ggplot2 [25].

### Siderophore production and detection

Bacterial strains were incubated on BM blood agar plates for 24 h to 48 h at 37°C. Subsequently, bacterial material was scratched from the plates, washed once with E-BHI medium, adjusted to an optical density at 600 nm (OD_600_) of 0.05 and incubated in 1 ml E-BHI for 72 h at 37°C and constant shaking at 160 rpm in 24 well plates. In case of *Corynebacterium accolens*, *Corynebacterium tuberculostearicum*, *Corynebacterium aurimucosum* and *Corynebacterium kroppenstedtii*, the medium was additionally supplemented with 0.4% Tween80 (Sigma-Aldrich); *C. acnes and L. clevelandensis* were incubated in 1.5-ml reaction tubes to minimize gas exchange with atmospheric oxygen; *Dolosigranulum pigrum* was incubated with 10% sterile-filtered spent medium of *C. accolens* to enhance growth [26]. After 72 h of incubation, the OD_600_ was determined, the bacterial cultures were centrifuged, and the supernatants were sterile filtered using 0.22 µm filter (Millex) to obtain the spent media. In order to quantify siderophore concentrations in the spent media, a siderophore detection kit (SideroTec Assay^TM^ from Emergen Bio) was used according to the manufacturer’s instructions with small alterations; instead of 100 µl samples, 50 µl samples were used and mixed with 50 µl Chelex treated MilliQ-purified H_2_O in order to reduce background signals of the medium.

### Analysis of growth support for S. *lugdunensis* under iron-restriction by xenosiderophores produced by nasal bacteria

In order to generate iron-restricted BHI agar plates, 2x BHI was treated with Chelex 100 resin (Sigma-Aldrich) for 24 h at 4°C and supplemented with 40 µM EDDHA. After sterile filtration, the medium was heated to 50°C and mixed 50:50 with 2x agar (50°C) containing 20% complement-inactivated horse serum (10% final concentration) (Sigma-Aldrich) to provide an iron source with complexed iron. Freshly grown *S. lugdunensis* was picked from blood agar plates, washed once with PBS, adjusted to an OD_600_ of 0.5, and streaked evenly onto the BHI agar plates. To evaluate the growth support for *S. lugdunensis* by the siderophore-containing spent media, 10 µl aliquots were spotted on previously plated *S. lugdunensis* cells as follows. Dependent on the endpoint OD_600_ of each culture, different volumes of spent medium were taken and mixed, if necessary, with fresh E-BHI to a final volume of 10 µl (for instance, from a culture with an endpoint OD_600_ of 0.2, 10 µl spent medium was taken whereas from a culture with an endpoint OD_600_ of 2.0, 1 µl spent medium was taken and mixed with 9 µl E-BHI medium). After spotting 10 µl of each spent medium, the plates were incubated for 20 h at 37°C. If the spent media contained siderophores that could promote growth of *S. lugdunensis*, zones of enhanced growth appeared that were characterized according to their diameters.

### Generation and complementation of S. lugdunensis and Staphylococcus epidermidis mutants

For the construction of *S. lugdunensis* and *S. epidermidis* mutants the thermosensitive plasmid pIMAY [27] and the primers listed in Suppl. Table 4 were used. Targeted mutagenesis was performed as described in [27]. In brief, 500 bp DNA fragments up- and downstream of the ferric hydroxamate uptake (*fhu*) gene locus for *S. lugdunensis* and of the staphyloferrin A biosynthetic genes (*sfaDAB*) for *S. epidermidis* were amplified by PCR using the corresponding primers ((1)/(2) and (3)/(4) for *fhu* knockout, (9)/(10) and (11)/(12) for *sfaDAB* knockout). The resulting PCR products contained overlapping sequences, which facilitated hybridization in order to construct the gene deletion sequence. This DNA sequence was inserted into pIMAY, digested via SacI/KpnI (Thermo) for *fhu* knockout and via SalI/SacI (Thermo) for *sfaDAB* knockout, using Gibson assembly (New England BioLabs) according to the manufacturer’s instructions. After transformation of *E. coli* DC10B and isolation of the plasmid construct, *S. lugdunensis* or *S. epidermidis* were transformed via electroporation and incubated at 30°C in TSB with 10 µg/ml chloramphenicol. Selection for plasmid integration into the chromosome was performed at 37°C in TSB with 10 µg/ml chloramphenicol, and positive integration clones were subsequently grown at 30°C without chloramphenicol to promote excision of the plasmid. The loss of plasmid was selected on TSA containing 1 µg/ml anhydrotetracycline and correct *S. lugdunensis Δfhu* or *S. epidermidis ΔsfaDAB* knock out clones were confirmed by chloramphenicol susceptibility testing and PCR (primers (5)/(6) for Δ*fhu*, (13)/(14) for Δ*sfaDAB)*.

For complementation in *S. lugdunensis*, primers (7)/(8) were used to amplify the wild type *fhu* gene. The DNA was inserted into SalI/XbaI digested pRB474 using Gibson assembly and transformed into *E. coli* DC10B. Subsequently, the complementation plasmid pRB474_*fhu* was used to transform *S. lugdunensis* Δ*fhu*. Chloramphenicol-resistant transformants were sub-cultured and correct complementation constructs were confirmed via PCR using primers (15)/(16).

## Results

### *S. lugdunensis* colonization reduces the risk of *S. aureus* carriage in healthy volunteers five-fold

We wondered if our recent report on a strong negative correlation between nasal colonization by *S. lugdunensis* and *S. aureus* [14] might have been confounded by underlying health problems or prior antibiotic treatment in the hospitalized patients included in this group. To assess the relation between *S. lugdunensis* and *S. aureus* in non-healthcare-associated humans a cohort of 270 healthy human volunteers was analyzed for nasal carriage of one or both bacterial species. The average age was 26.7 years and 63.7% of the participants were women. Nasal swabs were plated on selective media that allowed the specific detection of either *S. aureus* or *S. lugdunensis*. The identity of representative colonies from each donor was also verified by matrix-assisted laser desorption/ionization time-of-flight mass spectroscopy (MALDI-TOF).

The *S. aureus* carriage rate was 29.3%, which is very close to that of the hospitalized patient cohort and other previous studies [8, 14]. 6.3% of the participants were colonized by *S. lugdunensis* (Table 1), which is slightly lower than for the hospitalized patient cohort (9.1%). Nasal carriage of *S. lugdunensis* was much higher in male (12.3%) compared to female participants (2.9%). Notably, only one of the 270 participants was colonized by both, *S. aureus* and *S. lugdunensis*, which corresponds to a 5.2-fold lower propensity of *S. lugdunensis*-positive humans to carry *S. aureus* compared to *S. lugdunensis*-negative humans. This ratio corresponds well to the 5.9-fold reduced risk of *S. lugdunensis* carriers to be colonized by *S. aureus*, reported for the hospitalized patient’s cohort [14]. All *S. lugdunensis* isolates from healthy volunteers contained the lugdunin gene cluster and all *S. aureus* isolates were highly susceptible to lugdunin. Thus, the negative correlation between *S. lugdunensis* and *S. aureus* is the same in healthy and hospitalized human populations.

**Table 1.** *S. aureus* and *S. lugdunensis* distribution in study participants.

|  | *S. lugdunensis*-positive<br>(female/male) | *S. lugdunensis*-negative<br>(female/male) | Total<br>(female/male) |
| --- | --- | --- | --- |
| *S. aureus*-positive probands | 1 (0/1) | 78 (49/29) | 79 (49/30) |
| *S. aureus*-negative probands | 15 (5/10) | 176 (118/58) | 191 (123/68) |
| Total | 16 (5/11) | 254 (167/87) | 270 (172/98) |
| Risk of *S. aureus* carriage (%) | 5.9* (0.0/9.1) | 30.8* (29.3/33.3) | 29.3 (28.5/30.6) |

A chi-square test of independence was performed to examine the relation between gender and the ability to swim. The relation between these variables was significant, $\chi^2$ (1, N=270)=4.3503, \*p=.037. *S. lugdunensis*-positive probands are significantly less often colonized by *S. aureus* than *S. lugdunensis*-negative volunteers.

### *S. lugdunensis* colonization is more frequent than estimated and remains largely stable over time

To analyze the dynamics of nasal *S. lugdunensis* persistence or loss, the colonization status of eight carriers were monitored at four time points for a period of 23 months (probands A-H; Fig. 1A). In addition, four non-*S. lugdunensis* carriers were included as controls (probands I-L). One of the latter was a *S. aureus* carrier (proband I). In four participants (A-C, F) *S. lugdunensis* was detected at all four time points by cultivation on agar plates. In one carrier (G) *S. lugdunensis* was not detected after seven months but it reemerged after 18 and 23 months. In two (D, E) or one (H) carrier(s), *S. lugdunensis* could not be cultivated at two or three subsequent time points, respectively. Interestingly, *S. lugdunensis* could also be cultivated from the noses of all four initially negative control participants after seven months and, additionally, in the nose of one control participant (K) after 23 months (Fig. 1A). These data suggested that *S. lugdunensis* may have a pronounced capacity to persist in the noses of some humans while it temporarily emerges and disappears in others.

**Fig. 1.**
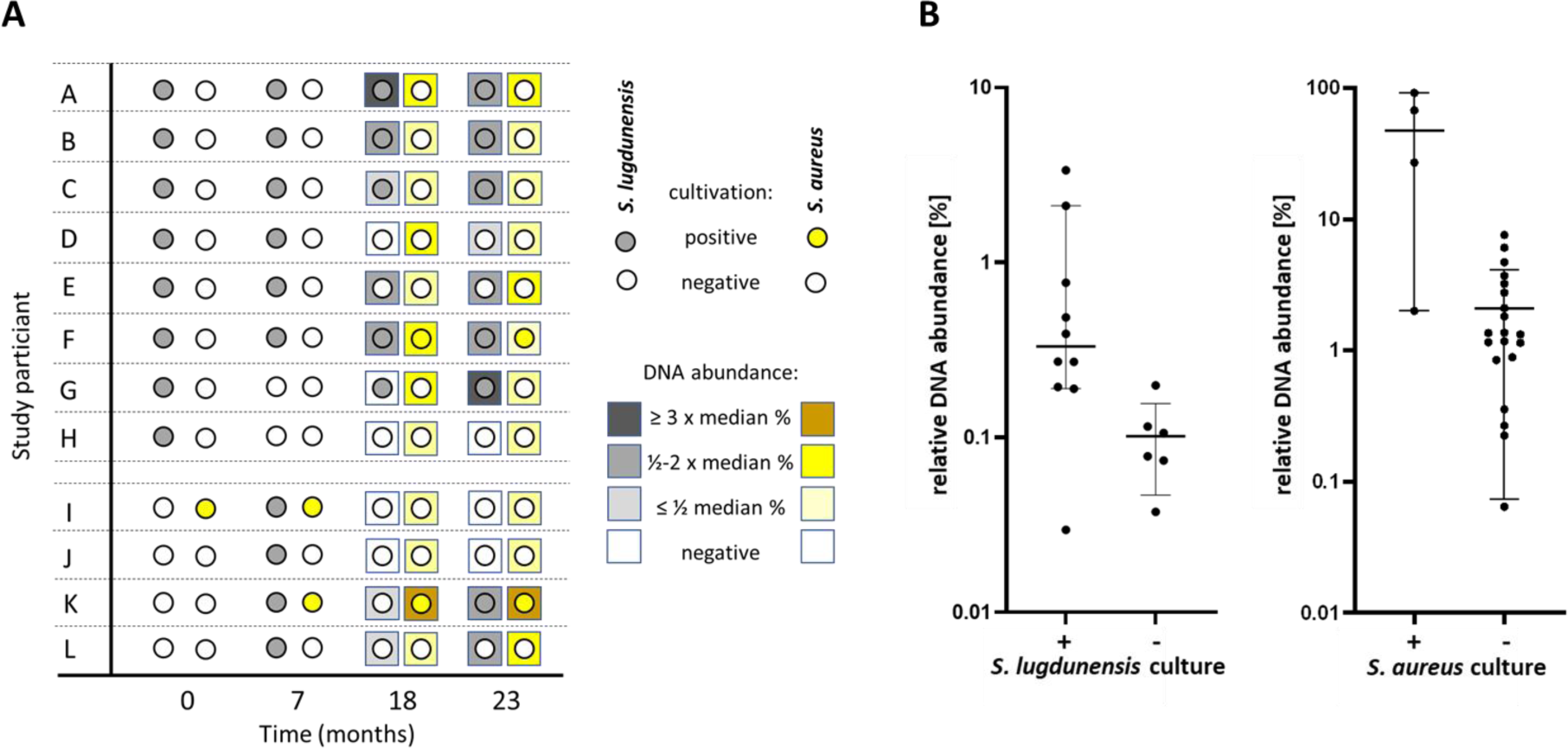
Detection of *S. lugdunensis* and *S. aureus* in the nose of study participants by cultivation or metagenome analysis. A) Culture outcomes for *S. lugdunensis* and *S. aureus* in the nasal samples and abundance of metagenome reads fort the two species. For each time point and each proband the culture results for *S. lugdunensis* (circles filled in grey) and *S. aureus* (circles filled in yellow) are shown (the control group of probands I – J, which was negative for S. lugdunensis at the first time point, is separated by two dashed lines). Negative culture outcome is indicated by empty circles. For time points 18 and 23 months, also the relative DNA abundances (=rel. proportions of reads specific for *S. lugdunensis* and *S. aureus*) in the metagenomes are indicated by colour-coded squares. The colour intensity indicates the relative DNA abundance compared to the corresponding median value of DNA abundance (see legend on the right). B) Relative proportions (in percent) of *S. lugdunensis* or *S. aureus* metagenome reads in microbiomes with positive or negative culture outcome. Left: relative proportions of *S. lugdunensis* reads versus culture outcome. Median of percentage for culture-positive samples: 0.33%; median of percentage for culture-negative samples: 0.09% (Mann-Whitney test, *p* = 0.0159). Right: relative proportions of *S. aureus* reads versus culture outcome. Median of percentage for culture-positive samples: 47.27%; median of percentage for culture-negative samples: 1.34% (Mann-Whitney test, p = 0.0072). Relative read proportions are shown on the y-axes in log scale. Medians are indicated by horizontal lines with 95% confidence intervals.

The nasal swabs were also analyzed for viable *S. aureus* cells. One of the *S. lugdunensis* carriers (F) became a *S. lugdunensis*-*S. aureus* co-carrier at months 18 and 23 (Fig. 1A). The initial *S. aureus* carrier from the control group (I) lost its *S. aureus* carrier status at months 18 and 23 while one of the initial non-carriers (K) became and remained positive after seven months. Co-carriage of *S. aureus* and *S. lugdunensis* was observed at individual time points in two participants (I, K) of the control group.

At two time points (18 and 23 months), the colonization status was also assessed by metagenome analysis of nasal swabs. To this end, bacterial isolates from nasal swabs were disintegrated, contaminating host DNA was degraded, and the remaining DNA was amplified and shotgun-sequenced on an Illumina system. An average of 30 million read pairs was obtained per sample, which were mapped to the NCBI nr protein database by DIAMOND and analyzed with the MEGAN6 algorithm. *S. lugdunensis* genome DNA was indeed found in most of the culture-positive and some additional samples (Fig. 1A) with a median relative abundance of *S. lugdunensis*-specific reads of 0.33% ranging from 0.03% to 3.37% (Suppl. Table 1). In only one of eleven culture-positive samples no *S. lugdunensis* DNA was found (Fig. 1A). Surprisingly, *S. lugdunensis* DNA was also detected in several of the culture-negative samples, at a low median relative abundance of 0.09% (range between 0.04% and 0.20%) indicating that *S. lugdunensis* may have remained present in most of the original carriers but, occasionally, at very low numbers, which did not reach the cultivation-dependent detection limit. Of the four original *S. lugdunensis* carriers who were culture-negative at one or more of the subsequent time points, three contained *S. lugdunensis* DNA even at culture-negative time points. Conversely, *S. lugdunensis* DNA was absent in the noses of two of the *S. lugdunensis* culture-negative control participants while two others contained some *S. lugdunensis* DNA with a relative abundance between 0.04% and 0.27% (the latter revealing a *S. lugdunensis*/*S. aureus* co-culture-positive state) (Fig. 1, Suppl. Table 1). Thus, *S. lugdunensis* is more frequent and colonizes human noses more consistently than estimated by swab cultivation. Samples with a positive culture outcome for *S. lugdunensis* revealed a significantly higher relative abundance of *S. lugdunensis* reads than those that were culture-negative (0.33% versus 0.09% median relative abundance, Mann-Whitney test, p=0.0159, Fig. 1B). Thus, the capacity to cultivate *S. lugdunensis* from carriers appears to depend on a threshold relative abundance in metagenomes, which probably lies somewhere between the median abundances observed for the cultivation-positive and cultivation-negative groups.

The metagenome analysis revealed similar findings for *S. aureus* as for *S. lugdunensis* - cultivation-positive samples had also high percentages of nasal *S. aureus* DNA with a median relative abundance of 47.3% (ranging from 2.0% to 91.4%) but even all culture-negative samples contained traces of *S. aureus* DNA (median relative abundance of 1.3%, ranging from 0.06% to 6.2%, Fig. 1A, Suppl. Table 1). These data suggest that *S. aureus* carriage is much higher in the human population than estimated via subcultivation of nasal swabs and that culture positivity depends on a threshold DNA abundance in metagenomes of above 2.0%. It is interesting to note that the threshold abundance for successful cultivation is obviously much higher for *S. aureus* than for *S. lugdunensis* (Fig. 1B).

The *S. lugdunensis* contigs from each consistently colonized proband (A, B, C, F and K) matched with specific S*. lugdunensis* genomes from databases of only one or two clonal complexes (CC1, CC3, CC6, CC7, or a currently undefined CC) per proband (Fig. 2) suggesting that a given human is usually colonized by one or only a few *S. lugdunensis* clones. This association remained largely the same at the two sampling times, pointing to a stable clonal distribution (samples G-2, K; see Fig. 2).

**Fig. 2.**
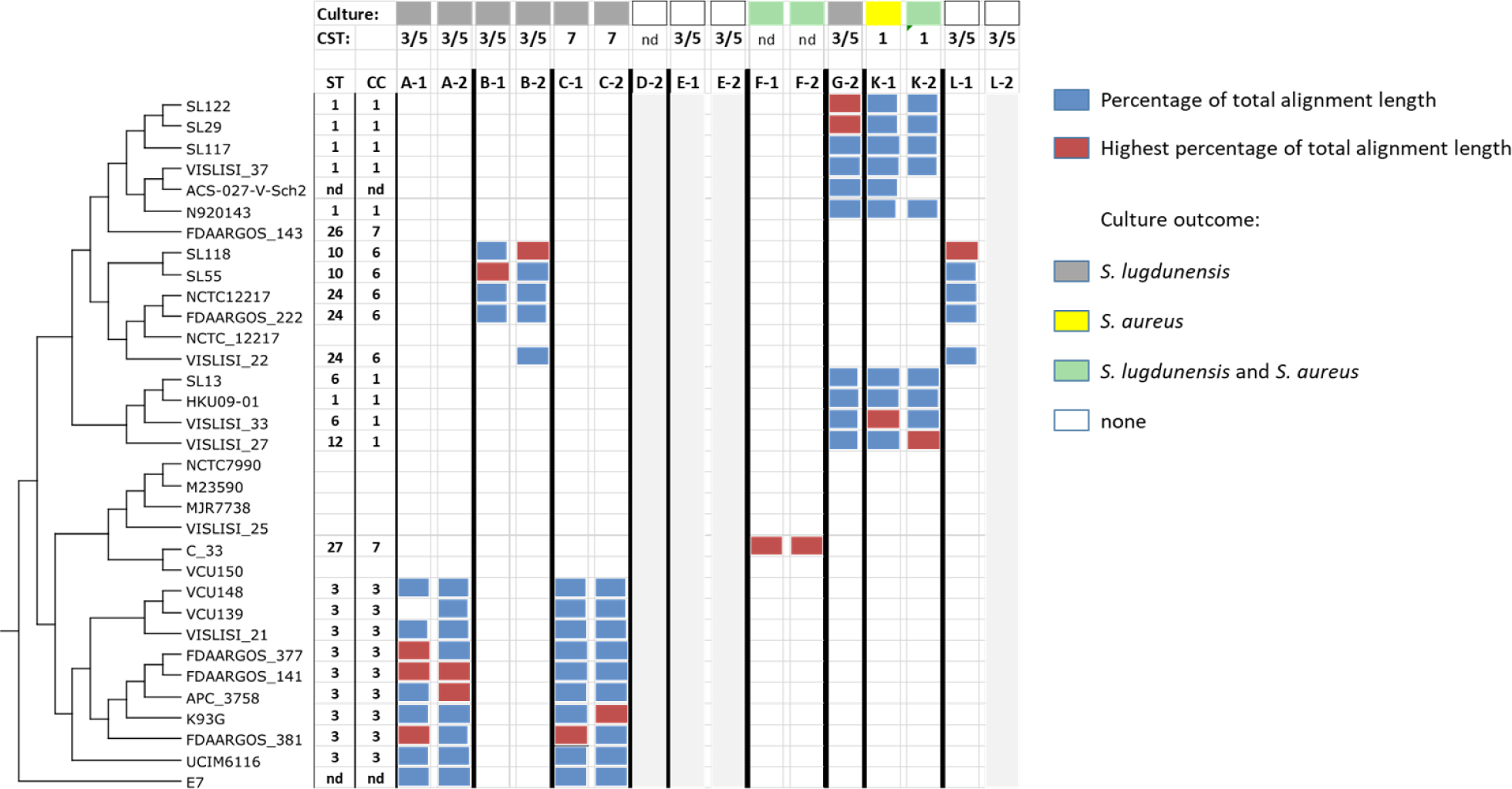
*S. lugdunensis*-containing metagenomes include only one or a few *S. lugdunensis* clones. *S. lugdunensis* strain profiles based on BLASTN comparisons of contigs from assembled metagenome reads with *S. lugdunensis* genome sequences. Sample numbers (1 or 2) indicate the corresponding metagenome analysis after 18 or 23 months, respectively. Blue bars represent the percentages (top 10%) of total hit alignment lengths between sample contigs and the corresponding *S. lugdunensis* strain (thresholds: 99.9% identity, 99.9% contig length in each hit alignment, only those contigs with a length of at least 250 nucleotides were considered). Red bars indicate highest percentage of total alignment length in the corresponding sample (see legend on top of right side). The phylogenetic tree of *S. lugdunensis* strains was downloaded from the NCBI database as Nexus-file and edited with Dendroscope [43]. Please note that genome data for *S. lugdunensis* NCTC12217 are presented twice in the NCBI tree as complete and as draft genome sequence. The strain profiling presented here is based on the complete genome sequence (NCTC12217). Volunteers who were tested positive for *S. lugdunensis* in the accompanying culture assays are indicated by grey rectangles, co-carriers by green rectangles, *S. aureus* carriers by yellow rectangles and non-carriers by non-filled rectangles (see legend on right side). Microbiomes of the CST3/5, CST1 and CST7 types are labelled accordingly. Samples labeled with areas shaded in light grey yielded no analyzable assemblies of the *S. lugdunensis* reads. Sequence types (ST) and corresponding clonal complexes (CC) of the profiled *S. lugdunensis* strains were obtained from https://bigsdb.web.pasteur.fr/staphlugdunensis. nd, no CST or ST/CC assignment possible.

### *S. lugdunensis* is associated with CSTs three, five, and seven

Based on the metagenome data the nasal microbiome composition of the two samples from twelve participants was elucidated and compared with the previously reported CST classification [11]. Notably, most of the metagenomes could be allocated to one of the seven CSTs (Fig.3A, Suppl. Table 1). Most S. *lugdunensis* carriers could be assigned to CST3 or CST5, which are dominated by *S. epidermidis* or *Corynebacterium sp.*, respectively, at both (participants A, B, E) or at least one time point (F, G, H). Several of these samples changed over time from CST5 to CST3 or vice versa. The two CSTs had very similar composition and clustered in principal component analysis (Fig. 3B). The dominance by either *S. epidermidis* or *Corynebacterium sp.* appears to vary over time, suggesting that CST3 and CST5 represent dynamic states of the same CST, which we refer to as CST3/5 from now on. Participant D had undergone systemic antibiotic therapy before the first microbiome sampling time point, which was probably the reason for the unusual metagenome composition, which changed from a non-CST-classifiable microbiome composition dominated by *Streptococcus* and *Staphylococcus* to an unprecedented nasal community dominated by *Haemophilus influenzeae*. The nasal microbiome of participant F was dominated by *Moraxella*, *Streptococcus*, and *Peptinophilus* and could also not be assigned to one of the established CSTs (Fig. 3A). *S. lugdunensis* associated most consistently with CST3/5, and all CST3/5 samples (except H-1) were found to contain *S. lugdunensis* genomic DNA at above-average abundances between 0.1% and 3.4% (median 0.3%) (Suppl. Table 1). The four control participants had CST7, CST1, or CST3/5, which were largely the same at the two different time points. Genomic DNA amounts in CST1, which is defined by *S. aureus* dominance, revealed very high *S. aureus* DNA abundances of 91.4 and 67.5%.

**Fig. 3.**
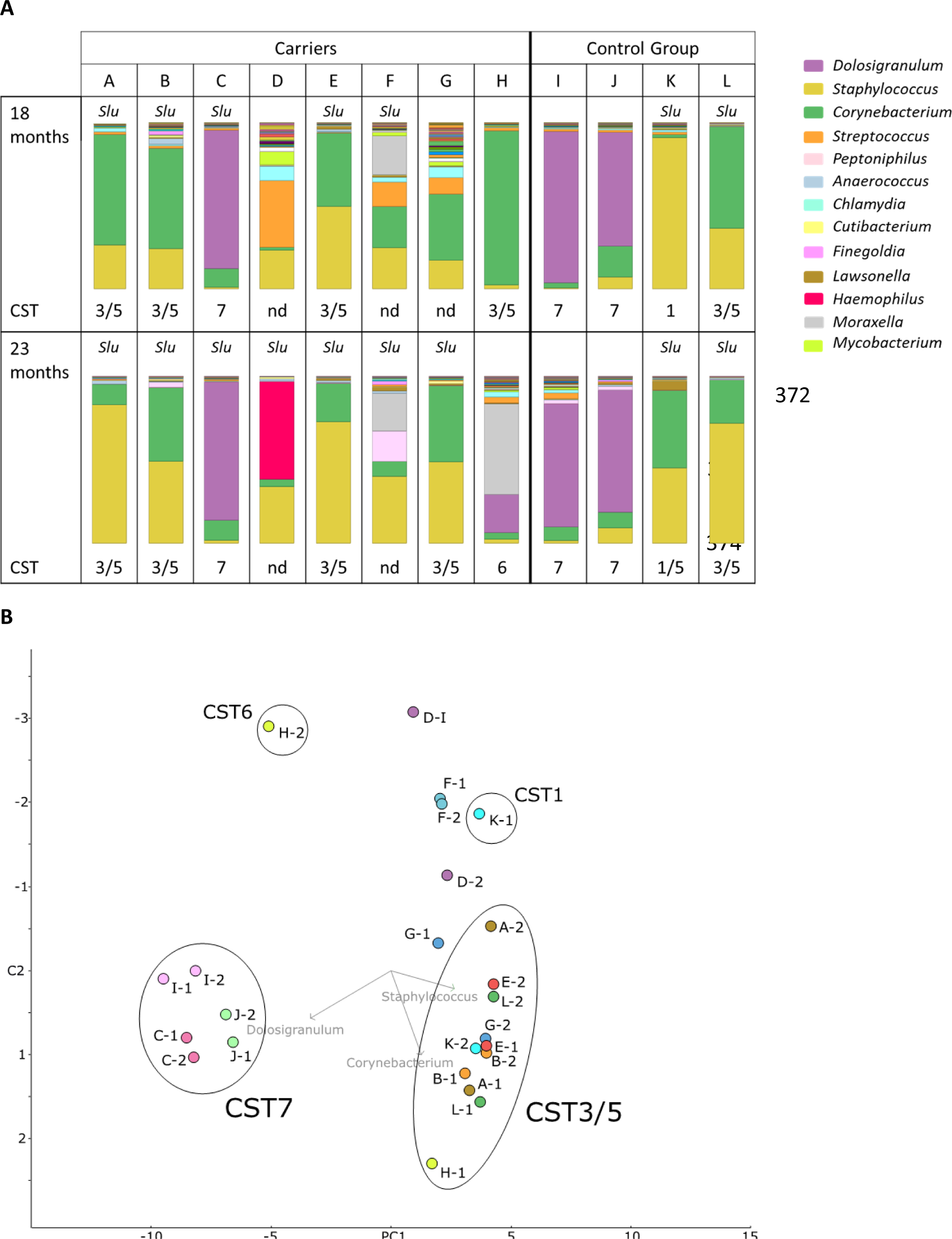
Most of *the S. lugdunensis*-colonized study participants can be assigned to CST3/5 and CST7. A) Nasal microbiome profiles at the genus level. Relative proportions of bacterial genera present in the nasal microbiomes at time points 18 months and 23 months are presented by stacked bar charts. The different bacterial genera are indicated by colours as shown in the legend on the right. The presence of *S. lugdunensis* reads in the metagenomes is indicated (*Slu*). The microbiome profiles that could be assigned to the community state types (CST) published by *Liu et al* [11] are labelled accordingly. Profiles not fitting to one of these types are designated “nd”. B) Clustering of the microbiome genus profiles by principal coordinates analysis. Comparison of the microbiome profiles at genus level by principal coordinates analysis based on evolutionary distances according to Hellinger [18]. The data showed the highest variation along principal component 1 (PC1, 52.6%). Principal component 2 (PC2) represents a variation of 15.0%. The microbiomes are represented by colored circles and numbers correlating with the corresponding metagenome analysis time point (-1, 18 months or -2, 23 months). Clusters representing the two main microbiome types CST3/5 and CST7 are labeled correspondingly. Vectors drawn and labeled in light grey point towards the prevalent genera in the clusters/subclusters. Note that sample K-2 clusters with the CST3/5 microbiomes but is *de facto* regarded as CST1 because the *Staphylococcus* proportion is mainly composed of *S. aureus*.

In addition to *S. epidermidis*, *S. lugdunensis*, and *S. aureus*, more than 50% of the metagenomes contained *Mammaliicoccus sciuri* (formerly *Staphylococcus sciuri*), albeit at low numbers, and at least one third also traces of other CoNS such as *Staphylococcus capitis* and *Staphylococcus caprae* (Suppl. Fig. 1). The microbiomes contained a remarkable diversity of *Corynebacterium* species but the abundance of specific species was similar for a specific CST (Suppl. Fig. 2). CST3/5 was dominated by *Corynebacterium accolens* with lower representation by *Corynebacterium tuberculosteraricum*, *Corynebacterium diphteriae,* and others (Tab. 2, Suppl. Fig. 2). In contrast, CST7 contained major amounts of genomic DNA from *Corynebacterium propinquum* and *Corynebacterium pseudodiphtheriticum* (Tab. 2, Suppl. Fig. 2). In conclusion, nasal *S. lugdunensis* carriage was associated with a high abundance of *S. epidermidis*, *C. accolens*, *C. tuberculostearicum* and, in one study participant, *D. pigrum*, *C. propinquum*, *C. pseudodiphtheriticum*.

**Table 2.**
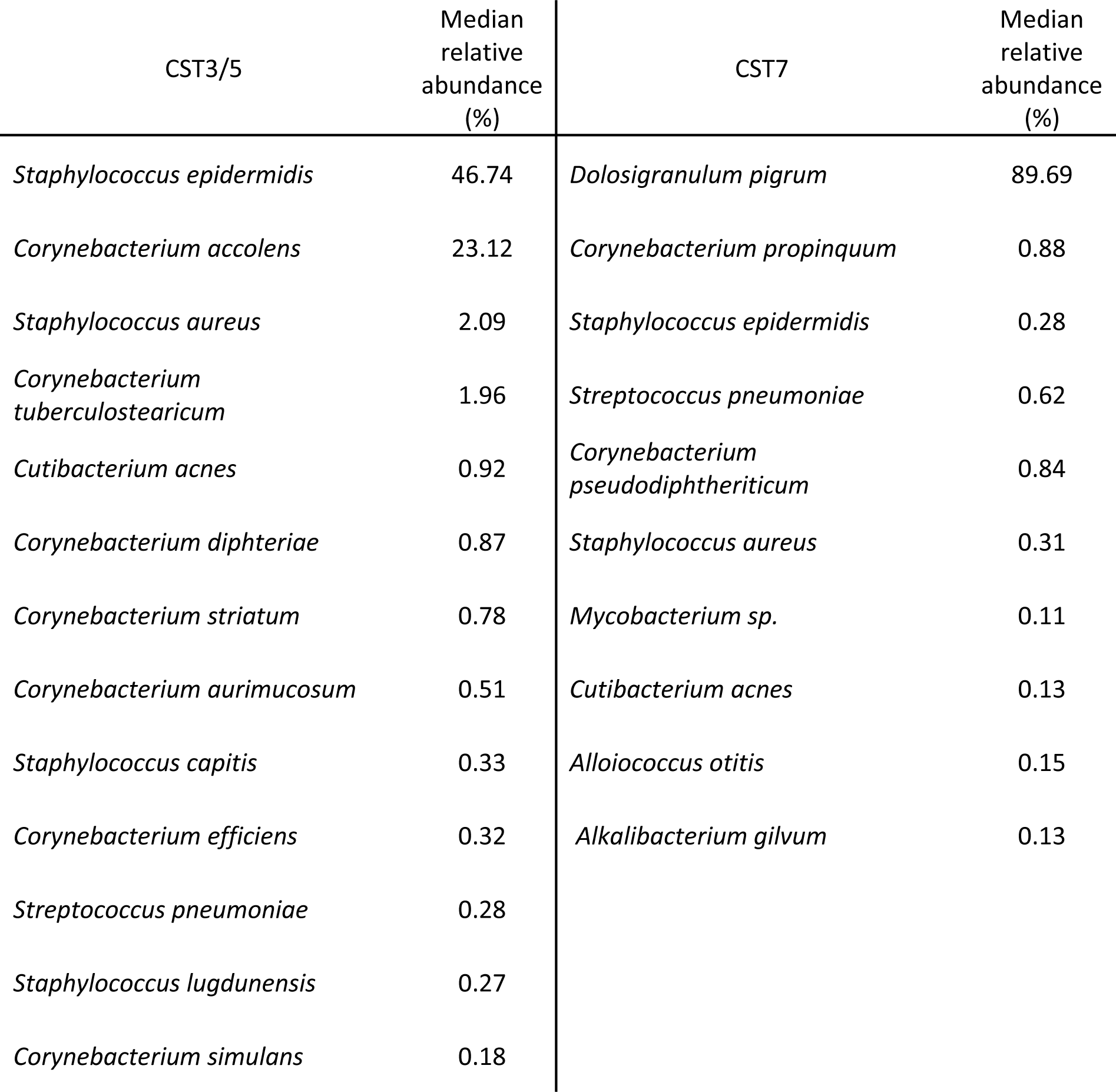
Bacterial species dominating the *S. lugdunensis*-associated CST3/5- and CST7-type microbiomes. Species that occur in all microbiomes of the indicated CST type are listed by decreasing median relative abundances (**%**) of species-specific metagenome reads. A threshold of 0.06% relative abundance was applied to the co-occurrence analysis with MEGAN6 a.

Quantitative metagenomic analysis permitted the assignment of bacterial species, which were negatively or positively correlated with the presence of *S. lugdunensis*. Based on a centered log-ratio-transformed abundance table of species reads we performed correlation network analyses with the Namap/Pearson correlation inference option of MetagenoNets [20] with thresholds for prevalence (0.1%) and occurrence (20%) (see Materials and Methods). Based on these parameters, we obtained correlation measures for 28 species throughout all 24 samples (Fig. 4, Suppl. Table 2). We observed correlation coefficients for *S. lugdunensis* ranging from -0.34 corresponding to the highest negative association with *Streptococcus pneumoniae* to +0.50, the strongest positive correlation with the *Clostridium*-related Bacillota (formerly Firmicutes) *Finegoldia magna* (Fig. 4). Most of the species dominating ST3/5 (Table 2) were also positively associated with *S. lugdunensis*. The data also revealed a strong negative association of *S. lugdunensis* with *S. aureus* (correlation coefficient -0.22) confirming our previous studies on the antagonistic interplay between these species [14] (Fig. 4). Moreover, several species, which are not typical nasal colonizers, found only at trace amounts were negatively associated with *S. lugdunensis*. In addition to the positive association with *Corynebacterium tuberculostearicum*, several other Actinobacteria including *Cutibacterium acnes*, *Corynebacterium striatum, Corynebacterium* efficiens, and the CoNS *S. epidermidis* and *S. capitis,* showed corresponding positive correlations with *S. lugdunensis* (Fig. 4. Suppl., Table 2).

**Fig. 4.**
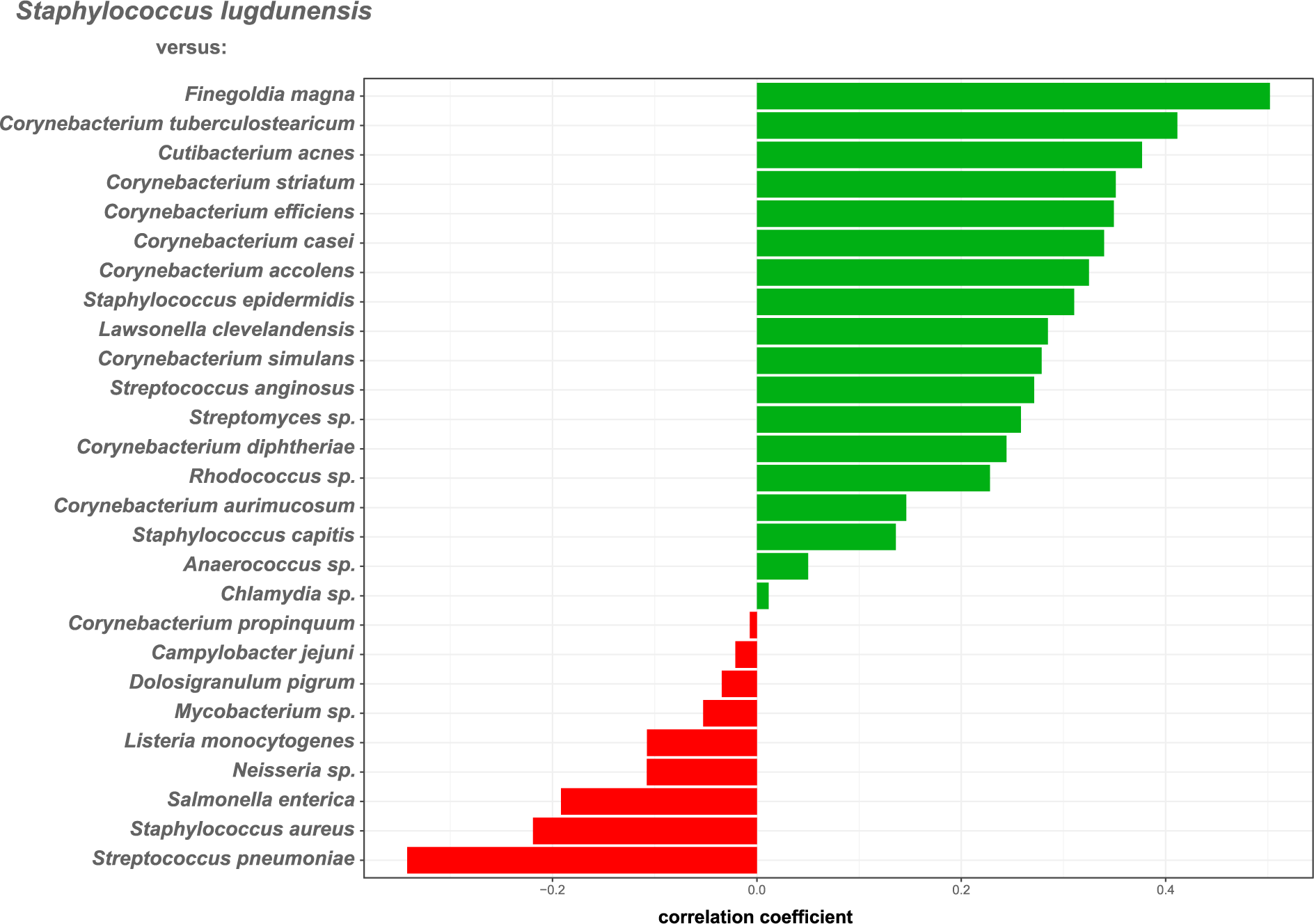
Specific bacterial species are positively or negatively associated with the abundance of *S. lugdunensis* reads in nasal metagenomes. Correlation coefficients were inferred based on a centered log-ratio transformed abundance table of species reads by correlation network analyses using the Namap/Pearson algorithm of MetagenoNets [20] with thresholds for prevalence and occurrence of 0.1% and 20%, respectively. Based on these parameters, we obtained correlation measures for 28 species throughout all 24 samples. Shown are the associations of *S. lugdunensis* with other microbiome species (shown at the x-axis). Positive associations are indicated by green bars, negative associations by red bars. The corresponding Pearson correlation coefficients are indicated at the x-axis.

The presence and abundance of *S. aureus* metagenome reads correlated negatively with *S. lugdunensis*, and other species, which have been shown to interfere with *S. aureus* colonization, including *D. pigrum*, *C. propinquum*, *S. pneumoniae*, and *F. magna* (Suppl. Fig. 3). In contrast, positive correlations for *S. aureus* were found for the Actinobacteria *Lawsonella clevelandensis*, *Corynebacterium aurimucosum*, the CoNS *S. epidermidis* and *S. capitis*, and the *Clostridium*-related Bacillota *Anaerococcus sp.*(Suppl. Fig. 3, Suppl. Table 2).

### Commensals dominating the *S. lugdunensis*-positive CST3/5 or CST7 support *S. lugdunensis* growth

The positive or negative association of *S. lugdunensis* with specific microbiome members raised the question, which mechanisms might be the basis for the antagonistic or mutualistic interactions. Some of the species among the typical nasal microbiome members, which were strongly negatively associated with *S. lugdunensis*, *S. pneumoniae* and *S. aureus* (Fig. 4), were highly susceptible to lugdunin (Fig. 5), which underscores the potential capacity of lugdunin to exclude specific microbiome members from the human nasal microbiome.

**Fig. 5.**
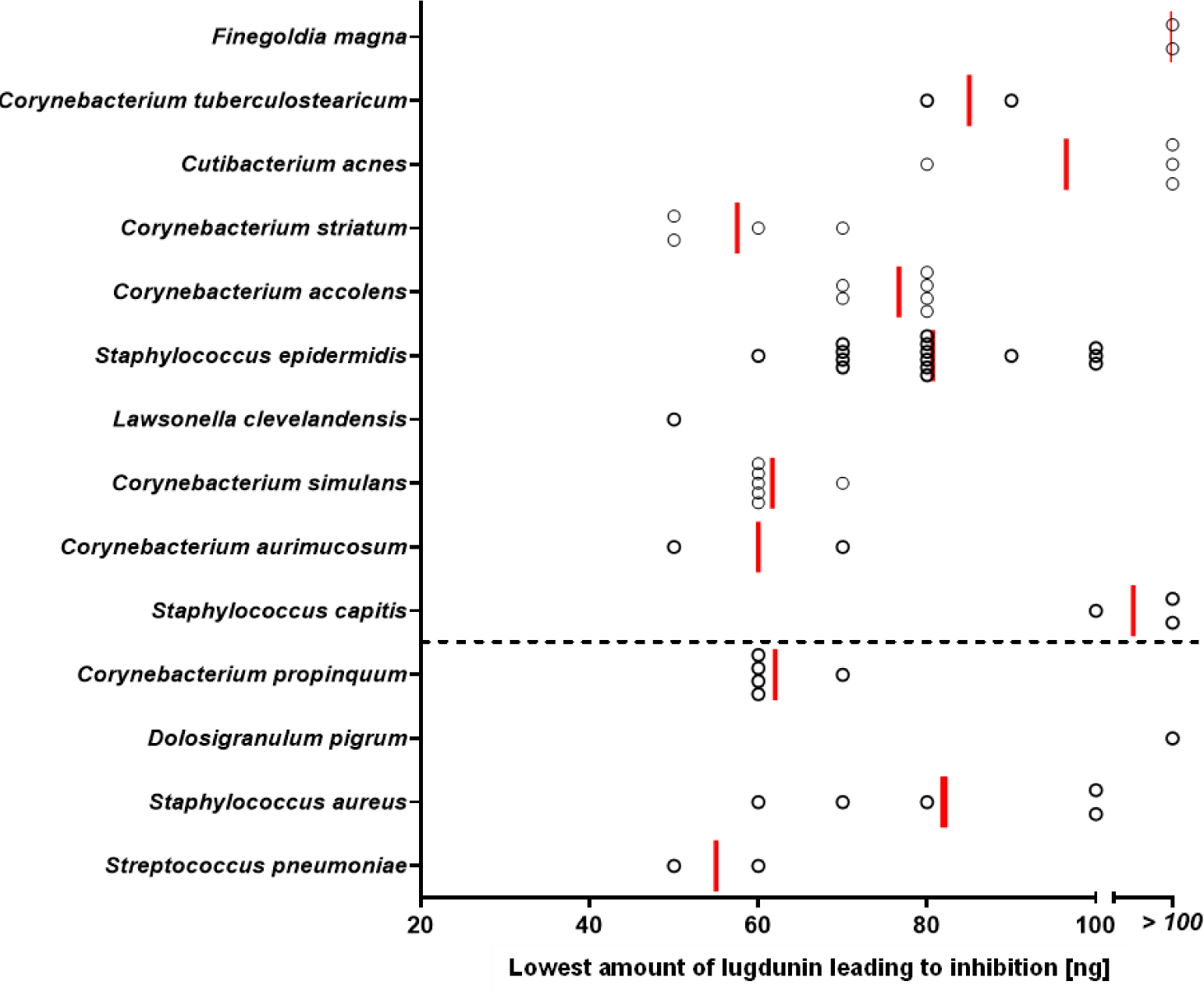
Lugdunin susceptibility of typical nasal bacterial species. Different amounts of lugdunin (0 to 100 ng in 2 μl) were spotted onto lawns of nasal bacteria, resulting in zones of inhibition after incubation for 24-48 h. Data points represent average values of three independent experiments, and vertical red lines show medians of each group. The dashed line indicates the threshold of positive (above) and negative (below) correlations of nasal bacteria with *S. lugdunensis*.

*S. lugdunensis* was positively associated with a particularly high number of nasal bacterial species (Fig. 4) raising the question how it can interact with so many different bacteria in a mutualistic manner. *S. lugdunensis* has been found to be strongly impaired in growth on iron-deprived media [28], and experimental data indicated that *S. lugdunensis* is probably unable to synthesize iron-sequestering siderophores and may therefore depend on uptake of siderophores produced by other nasal bacteria [29, 30]. This possibility was assessed by monitoring the growth of *S. lugdunensis* on iron-deficient agar plates supplemented with culture filtrates from other nasal bacteria. Culture filtrates of *C. tuberculostearicum*, *S. epidermidis*, and *S. capitis*, which were positively associated with *S. lugdunensis* (Fig. 4), had indeed a strong growth-promoting impact on *S. lugdunensis* (Fig. 6A). However, the tested isolates of a given bacterial species differed profoundly in their impact on *S. lugdunensis* growth indicating that the strength of mutualistic interaction with *S. lugdunensis* is probably strain specific.

**Fig. 6.**
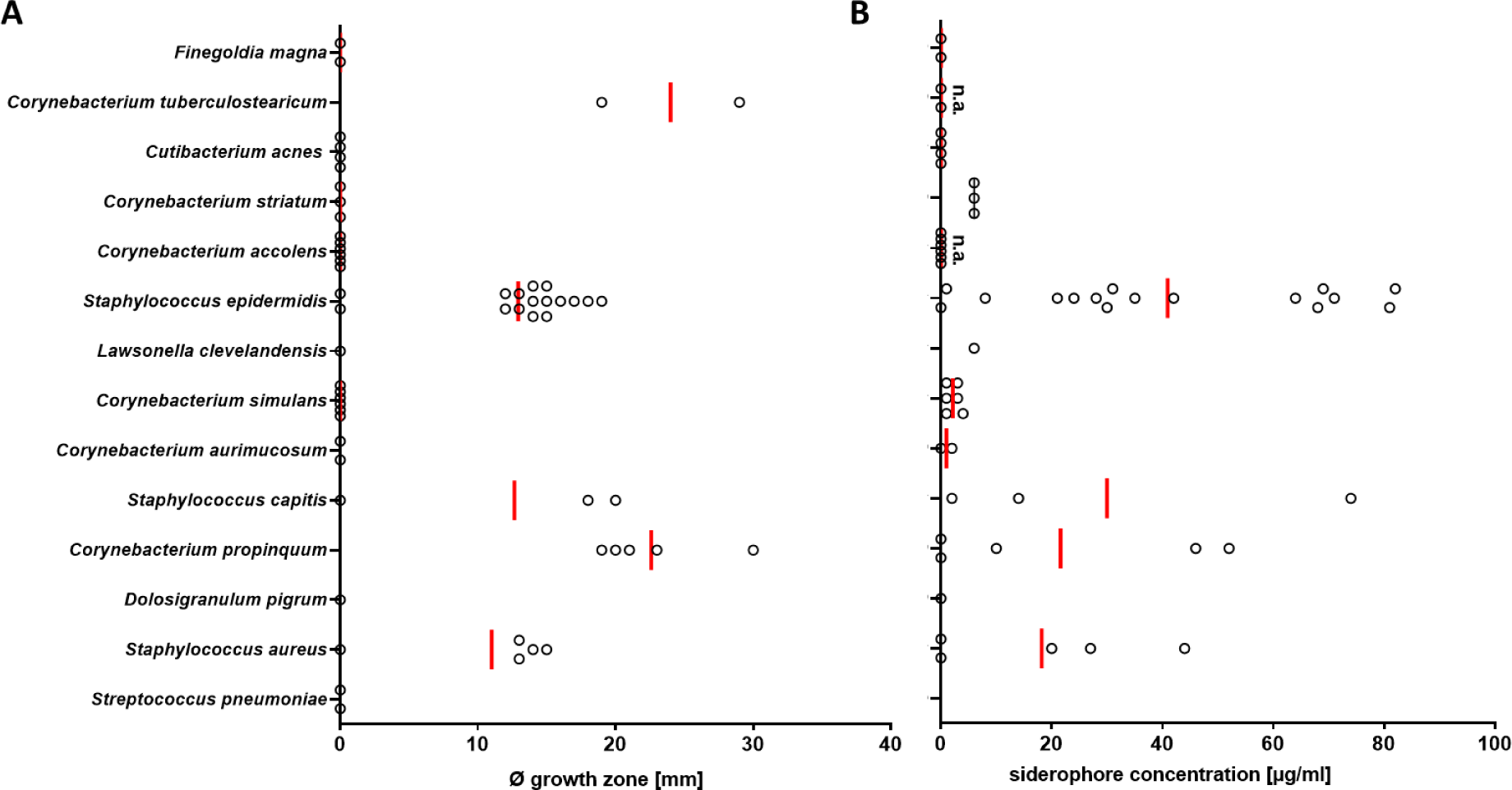
Nasal bacterial species producing siderophores have the capacity to support growth of *S. lugdunensis*. (A) Siderophore concentration in spent media from nasal bacteria after three days of incubation detected via SideroTec AssayTM kit. High concentrations of siderophores could be detected in spent media from staphylococcal species and *C. propinquum*; siderophore detection was not assessable for *C. accolens*, *C. aurimucosum*, and *C. tuberculostearicum* due to essential Tween 80 in growth medium that interacts with the reaction solutions provided by the SideroTec AssayTM kit. (B) Occurrence and sizes (diameter in mm) of growth zones of *S. lugdunensis* promoted by siderophore-containing spent media from nasal bacteria. Staphylococcal species, *C. propinquum* and *C. tuberculostearicum* support *S. lugdunensis* growth under iron-restricted conditions. Data points represent average values of three independent biological replicates, and vertical red lines show means of each species.

To assess if the growth-promoting bacterial microbiome members may release siderophores that *S. lugdunensis* can utilize, supernatants of cultures of the test bacteria grown under iron-restricted conditions were analysed for the presence of siderophores, by a colorimetric assay. Siderophore production was found, again, to be strongly strain-dependent, with some isolates of a given species showing no or very strong siderophore release (Fig. 6B). Strongly siderophore-producing isolates were found for *C. propinquum* and *S. epidermidis* but not for most of the nasal commensals that did not support growth of *S. lugdunensis*. *C. tuberculostearicum* could not be tested in this assay because essential components in its growth medium interfered with the colorimetric siderophore detection assay.

While *S. lugdunensis* lacks genes for siderophore synthesis, it encodes the siderophore uptake systems Hts and Sir allowing the acquisition of staphyloferrin A and B [29]. Additionally, it encodes the Fhu system recognizing hydroxamate-type and the Sst-system for acquisition of catechol/catecholamine type siderophores [31]. The *fhu* operon encoding also FhuC, the ATPase probably required for all of these uptake systems to function (except the SSt-system) [32] was deleted in *S. lugdunensis* HKU09-01 and IVK28 and the resulting mutants were compared to the wild type strains for their capacity to grow in the presence of siderophore-containing culture filtrates. The *fhu* mutants were unable to grow in the presence of culture filtrates from *C. propinquum* and *S. epidermidis* while the mutants complemented with plasmid-encoded copy of *fhu* grew equally well as the wild types (Fig. 7), which confirms the critical role of siderophore provision by other nasal bacteria for the capacity of *S. lugdunensis* to grow under conditions resembling the human nasal habitat.

**Fig. 7.**
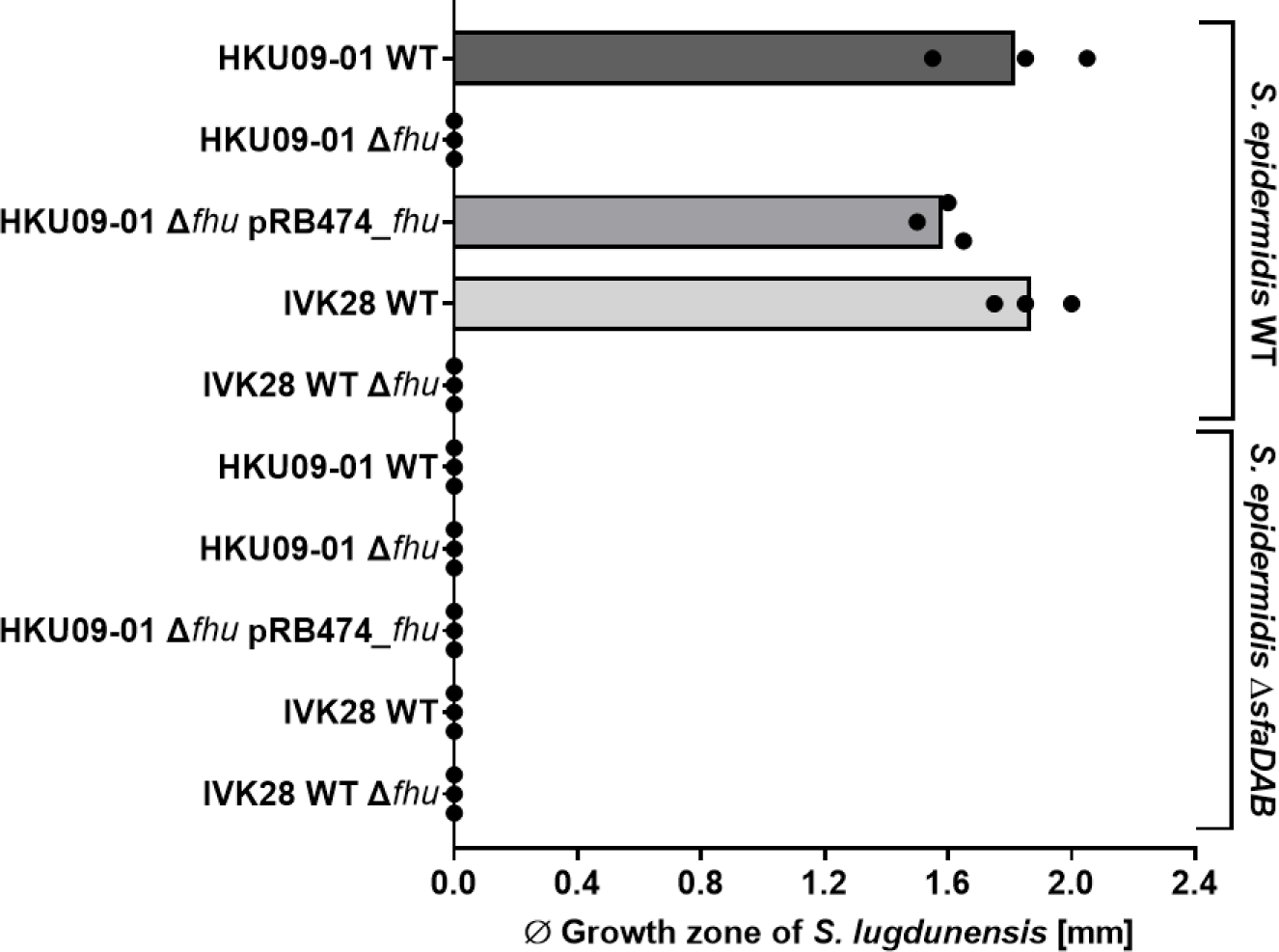
*S. lugdunensis* growth promotion requires the *fhu* siderophore uptake locus and depends on siderophore production by *S. epidermidis*. *S. lugdunensis* HKU09-01 and IVK28 wild-type (WT), isogenic Δ*fhu* mutants and complemented strains were grown on iron-restricted agar. Growth promotion was observed only for the wild-type and the complemented strains when spent media were spotted/added that contained staphyloferrin A produced by *S. epidermidis* WT. Bars represent the mean of three independent experiments and black dots values of single experiments.

## Discussion

The human nasal microbiome is less complex in composition than the intestinal microbiome [11]. Why the various human CSTs are dominated by specific bacterial species and why other species are found only in certain CSTs but absent from others has remained largely unclear. Nevertheless, a recent twin-cohort study has revealed that host genetics are of minor importance while the interaction among established microbiome residents appear to play key roles [11]. Our study demonstrates that even bacterial species that can be cultivated only from a minority of human nasal microbiomes are more prevalent than previously thought when analyses are based on metagenome data. Both, *S. lugdunensis* and *S. aureus* were identified in several nasal metagenomes from noses, which were culture-negative for these species. These data corroborate previous reports on the frequent presence of *S. aureus* DNA in nasal samples that did not yield positive *S. aureus* cultures [11, 33]. They also shed new light on the human *S. aureus* carrier status, which is obviously higher than the 30-40% previously estimated in culture-based studies. We found that the probability to cultivate *S. aureus* increased with the numbers of *S. aureus* reads in metagenomes indicating that a certain threshold number of *S. aureus* cells is necessary to achieve a positive culture. Bacterial numbers may change over time, which might explain the volatile *S. aureus* intermittent carrier state, characterized by a repeated change between culture-negative and culture-positive conditions with only low *S. aureus* counts per nare [34]. Very low numbers of *S. aureus* may not increase the risk of an at-risk patient to develop an invasive *S. aureus* infection. However, even very low *S. aureus* numbers may be of high relevance if it is an MRSA strain that might spread to other patients on a hospital ward and may elicit MRSA outbreaks. Future pathogen monitoring approaches should therefore not only rely on bacterial cultures but also on the analysis of metagenome data to identify even low numbers of notorious pathogens and their antibiotic resistance genes, for which bioinformatic tools have already been developed [35].

Many bacterial species, including *S. aureus* and *S. pneumoniae* were negatively associated with *S. lugdunensis* and were also highly susceptible to lugdunin. However, several species, which were positively associated with *S. lugdunensis*, including several coynebacterial and CoNS species, were also found to be lugdunin-susceptible, indicating that further criteria beyond the mere susceptibility to lugdunin need to be considered to explain the antagonistic interactions of *S. lugdunensis*. These might include the proximity or distance of *S. lugdunensis* to other microbiome members in a specific nasal sub-niche. In addition, lugdunin is also able to induce human-derived antimicrobial peptides which synergistically act especially against *S. aureus* [36]. It should be noted that the bacterial strains tested for susceptibility to lugdunin were representative isolates obtained from available strain collections, which might differ from strains in the nasal microbiomes of the study participants in their properties.

Several studies have been published with a focus on the bacterial competition in the human anterior nares [37–39], which found in part similar positive or negative associations as in our study. The results of our correlation analyses confirm previously described associations between *Corynebacterium*, *Staphylococcus*, *Cutibacterium, Dolosigranulum,* and *Streptococcus* species in the anterior nares [39]. We observed a negative association of *C. accolens* with *S. pneumoniae* (correlation coefficient -0.28), which corresponds to the reported antagonism of these two species [37]. Furthermore, we confirm the positive and negative associations of *D. pigrum* with other nasal microbiome species [38]. *D. pigrum* exhibited strong positive association with *C. propinquum* (correlation coefficient +0.66) and negative association with *S. aureus* (correlation coefficient -0.41) (Suppl. Table 2).

With microbiome precision editing approaches, it could become possible to eliminate *S. aureus* from the noses of at-risk patients or at least to reduce its numbers strongly enough to keep infection risks to a minimum [8]. Finding a way to establish robust nasal *S. lugdunensis* colonization in at-risk patients could become a sustainable way of preventing *S. aureus* carriage and infections. We provide strong evidence that *S. lugdunensis* is auxotrophic for iron-scavenging siderophores and may need other microbiome members releasing siderophores. Such compounds do not only support the producer but act as “public goods” for other bacteria with cognate siderophore uptake systems. Of note, the nutritional surrounding of the nasal cavity is iron-restricted and iron acquisition systems of *S. aureus* are expressed during colonization of humans and cotton rats [40, 41]. Additionally, we show in a co-submitted manuscript that siderophore acquisition of *S. aureus* is important during nasal colonization of cotton rats and that a plethora of siderophore-mediated interactions exist between nasal commensals and *S. aureus* (Zhao Y. et al – submitted)

*S. lugdunensis* does not produce siderophores but encodes ABC importers for carboxylate, hydroxamate, and catechol/catecholamine siderophores [29, 31]. Interestingly, *C. propinquum*, which supported *S. lugdunensis* growth under iron-limiting conditions, has been reported to bear genes for biosynthesis of the siderophore dehydroxynocardamine [42]. The capacities of different isolates of *S. epidermidis*, *S. hominis*, *S. capitis*, *S. aureus* to support *S. lugdunensis* growth was highly strain-dependent, which may point to substantial differences in the presence of such genes or their expression levels.

Several of the nasal bacterial species that correlated positively with *S. lugdunensis* did not support growth of *S. lugdunensis* under iron-restricted conditions. It is possible that the nasal strains differed in the production of siderophores from the test strains from our culture collection, used for in-vitro co-cultivation. On the other hand, some of the positive associations may result from higher-order interactions in larger bacterial networks with community members supporting other bacteria that were able to secrete siderophores.

## Conclusion

We demonstrate that the nasal cavity of only a minority of the human population is colonized by *S. lugdunensis*, which eliminates *S. aureus* by production of antimicrobial lugdunin. *S. lugdunensis* cannot produce siderophores, and it prefers nasal microbiomes dominated by bacterial species that support iron acquisition by *S. lugdunensis* by production of suitable siderophores. Understanding and harnessing such complex mechanism will allow the development of innovative strategies for pathogen exclusion form human microbiomes. Engineering *S. lugdunensis* to produce its own siderophores or co-administration of other commensals producing siderophores for *S. lugdunensis* could become valuable strategies for sustained prevention of *S. aureus* infections.

## Declarations

### Ethics approval and consent to participate

This study was approved by the Institutional Review Board for Human Subjects at the University of Tübingen (project number 577/2015A), and informed written consent was obtained from all study participants before sample collection.

### Consent for publication

Not applicable

### Availability of data and material

The dataset supporting the conclusions of this article can be retrieved from NCBI under the BioProject number PRJNA1078731.

### Authors’ contributions

Conceptualization and funding acquisition: R.R., B.O.T.S., and C.S. performed and evaluated the majority of experiments; R.R. assembled metagenomes and performed bioinformatic analysis. B.O.T.S. constructed the Δ*fhuC* strain for siderophore uptake experiments. R.R., S.H., B.K., and A.P. designed the study; A.P. acquired funding and wrote the first draft of the manuscript.

## Acknowledgements

The authors thank Vera Augsburger, Cosima Hirt, and Darya Belikova for excellent technical support and Stephanie Grond for synthetic lugdunin.

## References

1. Kost C, Patil KR, Friedman J, Garcia SL, Ralser M. Metabolic exchanges are ubiquitous in natural microbial communities. Nature Microbiology. 2023;8(12):2244–52; doi: 10.1038/s41564-023-01511-x.

2. Heilbronner S, Krismer B, Brötz-Oesterhelt H, Peschel A. The microbiome-shaping roles of bacteriocins. Nat Rev Microbiol. 2021;19(11):726–39; doi: 10.1038/s41579-021-00569-w.

3. Keith JW, Pamer EG. Enlisting commensal microbes to resist antibiotic-resistant pathogens. J Exp Med. 2019;216(1):10–9; doi: 10.1084/jem.20180399.

4. Tacconelli E, Sifakis F, Harbarth S, Schrijver R, van Mourik M, Voss A, et al. Surveillance for control of antimicrobial resistance. Lancet Infect Dis. 2018;18(3):e99–e106; doi: 10.1016/s1473-3099(17)30485-1.

5. Tacconelli E, Autenrieth IB, Peschel A. Fighting the enemy within. Science. 2017;355(6326):689-90; doi: 10.1126/science.aam6372.

6. Tsolis RM, Bäumler AJ. Gastrointestinal host-pathogen interaction in the age of microbiome research. Curr Opin Microbiol. 2020;53:78–89; doi: 10.1016/j.mib.2020.03.002.

7. Zhang ZJ, Lehmann CJ, Cole CG, Pamer EG. Translating Microbiome Research From and To the Clinic. Annu Rev Microbiol. 2022; doi: 10.1146/annurev-micro-041020-022206.

8. Krismer B, Weidenmaier C, Zipperer A, Peschel A. The commensal lifestyle of Staphylococcus aureus and its interactions with the nasal microbiota. Nat Rev Microbiol. 2017;15(11):675–87; doi: 10.1038/nrmicro.2017.104.

9. Lee AS, de Lencastre H, Garau J, Kluytmans J, Malhotra-Kumar S, Peschel A, Harbarth S. Methicillin-resistant Staphylococcus aureus. Nat Rev Dis Primers. 2018;4:18033; doi: 10.1038/nrdp.2018.33.

10. van Dalen R, Elsherbini AMA, Harms M, Alber S, Stemmler R, Peschel A. Secretory IgA impacts the microbiota density in the human nose. Microbiome. 2023;11(1):233; doi: 10.1186/s40168-023-01675-y.

11. Liu CM, Price LB, Hungate BA, Abraham AG, Larsen LA, Christensen K, et al. Staphylococcus aureus and the ecology of the nasal microbiome. Sci Adv. 2015;1(5):e1400216; doi: 10.1126/sciadv.1400216.

12. Janek D, Zipperer A, Kulik A, Krismer B, Peschel A. High Frequency and Diversity of Antimicrobial Activities Produced by Nasal Staphylococcus Strains against Bacterial Competitors. PLoS Pathog. 2016;12(8):e1005812; doi: 10.1371/journal.ppat.1005812.

13. Torres Salazar BO, Heilbronner S, Peschel A, Krismer B. Secondary Metabolites Governing Microbiome Interaction of Staphylococcal Pathogens and Commensals. Microb Physiol. 2021;31(3):198–216; doi: 10.1159/000517082.

14. Zipperer A, Konnerth MC, Laux C, Berscheid A, Janek D, Weidenmaier C, et al. Human commensals producing a novel antibiotic impair pathogen colonization. Nature. 2016;535(7613):511-6; doi: 10.1038/nature18634.

15. Ho PL, Leung SM, Tse H, Chow KH, Cheng VC, Que TL. Novel selective medium for isolation of Staphylococcus lugdunensis from wound specimens. J Clin Microbiol. 2014;52(7):2633–6; doi: 10.1128/JCM.00706-14.

16. Buchfink B, Xie C, Huson DH. Fast and sensitive protein alignment using DIAMOND. Nat Methods. 2015;12(1):59–60; doi: 10.1038/nmeth.3176.

17. Huson DH, Beier S, Flade I, Gorska A, El-Hadidi M, Mitra S, et al. MEGAN Community Edition - Interactive Exploration and Analysis of Large-Scale Microbiome Sequencing Data. PLoS Comput Biol. 2016;12(6):e1004957; doi: 10.1371/journal.pcbi.1004957.

18. Mitra S, Gilbert JA, Field D, Huson DH. Comparison of multiple metagenomes using phylogenetic networks based on ecological indices. ISME J. 2010;4(10):1236–42; doi: 10.1038/ismej.2010.51.

19. Nurk S, Meleshko D, Korobeynikov A, Pevzner PA. metaSPAdes: a new versatile metagenomic assembler. Genome Res. 2017;27(5):824–34; doi: 10.1101/gr.213959.116.

20. Nagpal S, Singh R, Yadav D, Mande SS. MetagenoNets: comprehensive inference and meta-insights for microbial correlation networks. Nucleic Acids Res. 2020;48(W1):W572–W9; doi: 10.1093/nar/gkaa254.

21. Faust K, Sathirapongsasuti JF, Izard J, Segata N, Gevers D, Raes J, Huttenhower C. Microbial co-occurrence relationships in the human microbiome. PLoS Comput Biol. 2012;8(7):e1002606; doi: 10.1371/journal.pcbi.1002606.

22. Yadav D, Ghosh TS, Mande SS. Global investigation of composition and interaction networks in gut microbiomes of individuals belonging to diverse geographies and age-groups. Gut Pathog. 2016;8:17; doi: 10.1186/s13099-016-0099-z.

23. Team RC: R: A Language and Environment for Statistical Computing. In.: R Foundation for Statistical Computing; 2020.

24. Cimentada MKaSJaJ: corrr: Correlations in R. In., R package version 0.4.2 edn; 2020.

25. Wickham H: ggplot2: Elegant Graphics for Data Analysis. In.: Springer-Verlag New York; 2016.

26. Brugger SD, Eslami SM, Pettigrew MM, Escapa IF, Henke MT, Kong Y, Lemon KP. Dolosigranulum pigrum cooperation and competition in human nasal microbiota. bioRxiv. 2020:678698; doi: 10.1101/678698.

27. Monk IR, Shah IM, Xu M, Tan MW, Foster TJ. Transforming the untransformable: application of direct transformation to manipulate genetically Staphylococcus aureus and Staphylococcus epidermidis. mBio. 2012;3(2); doi: 10.1128/mBio.00277-11.

28. Flannagan RS, Brozyna JR, Kumar B, Adolf LA, Power JJ, Heilbronner S, Heinrichs DE. In vivo growth of Staphylococcus lugdunensis is facilitated by the concerted function of heme and non-heme iron acquisition mechanisms. J Biol Chem. 2022;298(5):101823; doi: 10.1016/j.jbc.2022.101823.

29. Brozyna JR, Sheldon JR, Heinrichs DE. Growth promotion of the opportunistic human pathogen, Staphylococcus lugdunensis, by heme, hemoglobin, and coculture with Staphylococcus aureus. Microbiologyopen. 2014;3(2):182–95; doi: 10.1002/mbo3.162.

30. Sheldon JR, Heinrichs DE. The iron-regulated staphylococcal lipoproteins. Front Cell Infect Microbiol. 2012;2:41; doi: 10.3389/fcimb.2012.00041.

31. Lebeurre J, Dahyot S, Diene S, Paulay A, Aubourg M, Argemi X, et al. Comparative Genome Analysis of Staphylococcus lugdunensis Shows Clonal Complex-Dependent Diversity of the Putative Virulence Factor, ess/Type VII Locus. Front Microbiol. 2019;10:2479; doi: 10.3389/fmicb.2019.02479.

32. Sheldon JR, Heinrichs DE. Recent developments in understanding the iron acquisition strategies of gram positive pathogens. FEMS Microbiol Rev. 2015;39(4):592–630; doi: 10.1093/femsre/fuv009.

33. Cole AL, Sundar M, Lopez A, Forsman A, Yooseph S, Cole AM. Identification of Nasal Gammaproteobacteria with Potent Activity against Staphylococcus aureus: Novel Insights into the “Noncarrier” State. mSphere. 2021;6(1); doi: 10.1128/mSphere.01015-20.

34. van Belkum A, Verkaik NJ, de Vogel CP, Boelens HA, Verveer J, Nouwen JL, et al. Reclassification of Staphylococcus aureus nasal carriage types. J Infect Dis. 2009;199(12):1820–6; doi: 10.1086/599119.

35. Lanza VF, Baquero F, Martínez JL, Ramos-Ruíz R, González-Zorn B, Andremont A, et al. In-depth resistome analysis by targeted metagenomics. Microbiome. 2018;6; doi: ARTN 11 10.1186/s40168-017-0387-y.

36. Bitschar K, Sauer B, Focken J, Dehmer H, Moos S, Konnerth M, et al. Lugdunin amplifies innate immune responses in the skin in synergy with host- and microbiota-derived factors. Nat Commun. 2019;10(1):2730; doi: 10.1038/s41467-019-10646-7.

37. Bomar L, Brugger SD, Yost BH, Davies SS, Lemon KP. Corynebacterium accolens Releases Antipneumococcal Free Fatty Acids from Human Nostril and Skin Surface Triacylglycerols. mBio. 2016;7(1):e01725–15; doi: 10.1128/mBio.01725-15.

38. Brugger SD, Eslami SM, Pettigrew MM, Escapa IF, Henke MT, Kong Y, Lemon KP. Dolosigranulum pigrum Cooperation and Competition in Human Nasal Microbiota. mSphere. 2020;5(5); doi: 10.1128/mSphere.00852-20.

39. Hardy BL, Merrell DS. Friend or Foe: Interbacterial Competition in the Nasal Cavity. J Bacteriol. 2021;203(5); doi: 10.1128/JB.00480-20.

40. Burian M, Wolz C, Goerke C. Regulatory adaptation of Staphylococcus aureus during nasal colonization of humans. PLoS One. 2010;5(4):e10040; doi: 10.1371/journal.pone.0010040.

41. Burian M, Rautenberg M, Kohler T, Fritz M, Krismer B, Unger C, et al. Temporal expression of adhesion factors and activity of global regulators during establishment of Staphylococcus aureus nasal colonization. J Infect Dis. 2010;201(9):1414–21; doi: 10.1086/651619.

42. Stubbendieck RM, May DS, Chevrette MG, Temkin MI, Wendt-Pienkowski E, Cagnazzo J, et al. Competition among Nasal Bacteria Suggests a Role for Siderophore-Mediated Interactions in Shaping the Human Nasal Microbiota. Appl Environ Microbiol. 2019;85(10); doi: 10.1128/AEM.02406-18.

43. Huson DH, Scornavacca C. Dendroscope 3: an interactive tool for rooted phylogenetic trees and networks. Syst Biol. 2012;61(6):1061–7; doi: 10.1093/sysbio/sys062.

44. Kaspar U, Kriegeskorte A, Schubert T, Peters G, Rudack C, Pieper DH, et al. The culturome of the human nose habitats reveals individual bacterial fingerprint patterns. Environ Microbiol. 2016;18(7):2130–42; doi: 10.1111/1462-2920.12891.

45. Lemon KP: Personal communication. In.; 2018.

46. Diep BA, Otto M. The role of virulence determinants in community-associated MRSA pathogenesis. Trends Microbiol. 2008;16(8):361–9; doi: 10.1016/j.tim.2008.05.002.

47. Tse H, Tsoi HW, Leung SP, Lau SK, Woo PC, Yuen KY. Complete genome sequence of Staphylococcus lugdunensis strain HKU09-01. J Bacteriol. 2010;192(5):1471–2; doi: 10.1128/JB.01627-09.

48. Bruckner R. A series of shuttle vectors for Bacillus subtilis and Escherichia coli. Gene. 1992;122(1):187–92; doi: 10.1016/0378-1119(92)90048-t.

