## Supplemental Tables S1-S4 for "Siderophore piracy enables the nasal commensal *Staphylococcus lugdunensis* to antagonize the pathogen *Staphylococcus aureus*"

**Suppl. Table. 1.** Comparison of metagenome data with community state types (CST). Relative proportions (in percent) of selected taxa that had been described as signature for community state types (CST) as described in [11]. Most abundant taxa are highlighted in bold. Round 1 or 2 indicates the corresponding time of metagenome analysis after 18 months or 23 months, respectively. CST: classification of designated CST types according to the prevalence of indicator taxa in the microbiomes or none if not determinable (nd). CST6 for sample H-2 was based on a relative proportion of 54.4 % *Moraxella* (not listed). For *S. lugdunensis,* the corresponding relative DNA abundances are also shown (in percent). The culture-based carrier states determined in parallel to the metagenome sequencing are indicated under "Culture": Slu, *S. lugdunensis* carrier; Sau, *S. aureus* carrier; co, co-carrier; non, non-carrier.

| Sample | Round | *Corynebacterium* | *S. epidermidis* | *S. aureus* | *Dolosigranulum* | CST | *S. lugdunensis* | Culture |
|---|---|---|---|---|---|---|---|---|
| **A** | 1 | **67.2** | 21.6 | 3.7 | 0.0 | **3/5** | 2.11 | Slu |
| | 2 | 12.4 | **69.3** | 4.7 | 0.0 | **3/5** | 0.77 | Slu |
| **B** | 1 | **60.4** | 14.3 | 0.9 | 0.0 | **3/5** | 0.49 | Slu |
| | 2 | **44.3** | 37.7 | 1.4 | 0.0 | **3/5** | 0.27 | Slu |
| **C** | 1 | 11.2 | 0.2 | 0.2 | **83.3** | **7** | 0.03 | Slu |
| | 2 | 12.0 | 0.3 | 0.1 | **82.9** | **7** | 0.19 | Slu |
| **D** | 1 | 3.6 | 7.3 | 6.2 | 0.0 | nd | 0.00 | non |
| | 2 | 4.4 | 32.0 | 1.4 | 0.0 | nd | 0.07 | non |
| **E** | 1 | 44.1 | **51.3** | 2.1 | 0.0 | **3/5** | 0.12 | non |
| | 2 | 23.1 | **71.5** | 3.2 | 0.0 | **3/5** | 0.20 | non |
| **F** | 1 | 24.8 | 1.6 | 27.0 | 0.0 | nd | 0.19 | co |
| | 2 | 9.1 | 26.5 | 2.0 | 0.1 | nd | 0.39 | co |
| **G** | 1 | 39.8 | 7.3 | 6.1 | 0.0 | nd | 0.00 | Slu |
| | 2 | **45.6** | **46.7** | 1.3 | 0.0 | **3/5** | 3.37 | Slu |
| **H** | 1 | **92.9** | 2.0 | 1.2 | 0.0 | **3/5** | 0.00 | non |
| | 2 | 3.9 | 0.2 | 1.8 | 23.0 | **6** | 0.00 | non |
| **I** | 1 | 3.0 | 0.1 | 0.3 | **91.2** | **7** | 0.00 | non |
| | 2 | 8.2 | 0.1 | 1.2 | **73.7** | **7** | 0.00 | non |
| **J** | 1 | 18.6 | 2.2 | 0.8 | **68.6** | **7** | 0.00 | non |
| | 2 | 9.3 | 3.1 | 0.4 | **73.3** | **7** | 0.00 | non |
| **K** | 1 | 1.8 | 0.5 | **91.4** | 0.0 | **1** | 0.04 | Sau |
| | 2 | **46.5** | 1.3 | **67.5** | 0.1 | **1** | 0.27 | co |
| **L** | 1 | **61.2** | 44.6 | 1.2 | 0.0 | **3/5** | 0.08 | non |
| | 2 | 26.0 | **73.2** | 2.8 | 0.0 | **3/5** | 0.11 | non |

**Suppl. Table 2.** Correlation coefficients of species present in the nasal microbiomes. Correlation coefficients were inferred based on a centered log-ratio-transformed abundance table of species reads by correlation network analyses using the Namap/Pearson algorithm of MetagenoNets [20] with thresholds for prevalence and occurrence of 0.1% and 20%, respectively. The table shows the resulting correlation measures for 28 species throughout all 24 samples. Positive correlations are emphasized by green shades (with intensity corresponding to the hight of the values), while negative correlations are shaded in red.

| | *Anaerococcus sp.* | *Campylobacter jejuni* | *Chlamydia sp.* | *Corynebacterium accolens* | *Corynebacterium aurimucosum* | *Corynebacterium casei* | *Corynebacterium diphtheriae* | *Corynebacterium efficiens* | *Corynebacterium propinquum* | *Corynebacterium simulans* | *Corynebacterium striatum* | *Corynebacterium tuberculostearicum* | *Cutibacterium acnes* | *Dolosigranulum pigrum* | *Finegoldia magna* | *Lawsonella clevelandensis* | *Listeria monocytogenes* | *Mycobacterium sp.* | *Neisseria sp.* | *Rhodococcus sp.* | *Salmonella enterica* | *Staphylococcus aureus* | *Staphylococcus capitis* | *Staphylococcus epidermidis* | *Staphylococcus lugdunensis* | *Streptococcus anginosus* | *Streptococcus pneumoniae* | *Streptomyces sp.* |
|---|---|---|---|---|---|---|---|---|---|---|---|---|---|---|---|---|---|---|---|---|---|---|---|---|---|---|---|---|
| *Anaerococcus sp.* | 1.00 | 0.26 | -0.14 | 0.14 | 0.47 | 0.08 | 0.19 | 0.24 | -0.26 | 0.07 | 0.18 | 0.56 | 0.62 | -0.16 | 0.27 | 0.24 | 0.23 | -0.08 | 0.20 | 0.28 | 0.20 | 0.41 | 0.27 | 0.66 | 0.05 | 0.19 | -0.54 | 0.25 |
| *Campylobacter jejuni* | 0.26 | 1.00 | 0.43 | -0.02 | 0.42 | 0.09 | 0.27 | -0.09 | 0.14 | 0.08 | 0.18 | 0.46 | 0.29 | 0.15 | 0.08 | 0.05 | 0.67 | 0.23 | 0.96 | 0.15 | 0.93 | 0.26 | -0.09 | 0.15 | -0.02 | 0.17 | -0.03 | 0.14 |
| *Chlamydia sp.* | -0.14 | 0.43 | 1.00 | -0.34 | -0.21 | -0.36 | -0.23 | -0.19 | 0.13 | 0.12 | -0.15 | -0.05 | -0.21 | -0.01 | -0.05 | 0.09 | 0.32 | 0.43 | 0.42 | 0.01 | 0.42 | 0.06 | -0.27 | -0.28 | 0.01 | 0.00 | 0.36 | 0.13 |
| *Corynebacterium accolens* | 0.14 | -0.02 | -0.34 | 1.00 | 0.60 | 0.32 | 0.66 | 0.27 | -0.36 | 0.43 | 0.74 | 0.54 | 0.56 | -0.04 | 0.15 | 0.51 | 0.15 | 0.02 | -0.16 | 0.50 | -0.12 | 0.07 | 0.20 | 0.59 | 0.32 | 0.52 | -0.28 | 0.40 |
| *Corynebacterium aurimucosum* | 0.47 | 0.42 | -0.21 | 0.60 | 1.00 | 0.52 | 0.83 | 0.21 | -0.25 | 0.59 | 0.79 | 0.85 | 0.76 | -0.14 | 0.19 | 0.52 | 0.40 | 0.17 | 0.34 | 0.57 | 0.37 | 0.46 | 0.37 | 0.59 | 0.15 | 0.56 | -0.30 | 0.46 |
| *Corynebacterium casei* | 0.08 | 0.09 | -0.36 | 0.32 | 0.52 | 1.00 | 0.49 | 0.18 | 0.19 | 0.40 | 0.42 | 0.36 | 0.27 | 0.15 | 0.28 | 0.15 | -0.07 | -0.14 | 0.16 | 0.08 | 0.15 | 0.16 | 0.56 | 0.38 | 0.34 | 0.05 | -0.38 | -0.03 |
| *Corynebacterium diphtheriae* | 0.19 | 0.27 | -0.23 | 0.66 | 0.83 | 0.49 | 1.00 | 0.28 | -0.22 | 0.67 | 0.90 | 0.64 | 0.72 | -0.06 | 0.23 | 0.63 | 0.44 | 0.13 | 0.16 | 0.77 | 0.14 | 0.32 | 0.33 | 0.58 | 0.24 | 0.73 | -0.24 | 0.64 |
| *Corynebacterium efficiens* | 0.24 | -0.09 | -0.19 | 0.27 | 0.21 | 0.18 | 0.28 | 1.00 | -0.27 | 0.26 | 0.42 | 0.19 | 0.33 | 0.01 | 0.34 | 0.06 | 0.05 | -0.06 | -0.14 | 0.64 | -0.22 | -0.06 | 0.46 | 0.32 | 0.35 | 0.59 | -0.37 | 0.51 |
| *Corynebacterium propinquum* | -0.26 | 0.14 | 0.13 | -0.36 | -0.25 | 0.19 | -0.22 | -0.27 | 1.00 | 0.05 | -0.45 | -0.20 | -0.22 | 0.66 | -0.13 | -0.30 | 0.11 | -0.01 | 0.23 | -0.32 | 0.28 | -0.16 | -0.07 | -0.20 | -0.01 | -0.31 | 0.28 | -0.30 |
| *Corynebacterium simulans* | 0.07 | 0.08 | 0.12 | 0.43 | 0.59 | 0.40 | 0.67 | 0.26 | 0.05 | 1.00 | 0.66 | 0.46 | 0.50 | 0.13 | 0.04 | 0.45 | 0.35 | 0.53 | 0.00 | 0.67 | 0.08 | 0.31 | 0.41 | 0.31 | 0.28 | 0.65 | -0.01 | 0.65 |
| *Corynebacterium striatum* | 0.18 | 0.18 | -0.15 | 0.74 | 0.79 | 0.42 | 0.90 | 0.42 | -0.45 | 0.66 | 1.00 | 0.62 | 0.66 | -0.24 | 0.32 | 0.60 | 0.34 | 0.22 | 0.06 | 0.76 | 0.05 | 0.27 | 0.33 | 0.49 | 0.35 | 0.78 | -0.23 | 0.67 |
| *Corynebacterium tuberculostearicum* | 0.56 | 0.46 | -0.05 | 0.54 | 0.85 | 0.36 | 0.64 | 0.19 | -0.20 | 0.46 | 0.62 | 1.00 | 0.86 | -0.17 | 0.28 | 0.50 | 0.33 | 0.11 | 0.36 | 0.49 | 0.35 | 0.26 | 0.25 | 0.65 | 0.41 | 0.50 | -0.49 | 0.46 |
| *Cutibacterium acnes* | 0.62 | 0.29 | -0.21 | 0.56 | 0.76 | 0.27 | 0.72 | 0.33 | -0.22 | 0.50 | 0.66 | 0.86 | 1.00 | -0.11 | 0.27 | 0.52 | 0.35 | 0.07 | 0.14 | 0.66 | 0.13 | 0.27 | 0.28 | 0.74 | 0.38 | 0.62 | -0.54 | 0.59 |
| *Dolosigranulum pigrum* | -0.16 | 0.15 | -0.01 | -0.04 | -0.14 | 0.15 | -0.06 | 0.01 | 0.66 | 0.13 | -0.24 | -0.17 | -0.11 | 1.00 | -0.16 | -0.41 | 0.19 | -0.01 | 0.13 | -0.07 | 0.19 | -0.41 | -0.07 | -0.08 | -0.03 | -0.17 | 0.15 | -0.01 |
| *Finegoldia magna* | 0.27 | 0.08 | -0.05 | 0.15 | 0.19 | 0.28 | 0.23 | 0.34 | -0.13 | 0.04 | 0.32 | 0.28 | 0.27 | -0.16 | 1.00 | 0.33 | -0.18 | -0.38 | 0.07 | 0.22 | -0.08 | -0.10 | -0.10 | 0.28 | 0.50 | 0.20 | -0.41 | 0.30 |
| *Lawsonella clevelandensis* | 0.24 | 0.05 | 0.09 | 0.51 | 0.52 | 0.15 | 0.63 | 0.06 | -0.30 | 0.45 | 0.60 | 0.50 | 0.52 | -0.41 | 0.33 | 1.00 | 0.17 | -0.05 | -0.04 | 0.56 | -0.10 | 0.46 | 0.07 | 0.49 | 0.28 | 0.55 | -0.18 | 0.50 |
| *Listeria monocytogenes* | 0.23 | 0.67 | 0.32 | 0.15 | 0.40 | -0.07 | 0.44 | 0.05 | 0.11 | 0.35 | 0.34 | 0.33 | 0.35 | 0.19 | -0.18 | 0.17 | 1.00 | 0.54 | 0.55 | 0.47 | 0.58 | 0.32 | 0.13 | 0.21 | -0.11 | 0.44 | 0.25 | 0.42 |
| *Mycobacterium sp.* | -0.08 | 0.23 | 0.43 | 0.02 | 0.17 | -0.14 | 0.13 | -0.06 | -0.01 | 0.53 | 0.22 | 0.11 | 0.07 | -0.01 | -0.38 | -0.05 | 0.54 | 1.00 | 0.15 | 0.21 | 0.26 | 0.30 | 0.16 | -0.07 | -0.05 | 0.25 | 0.39 | 0.31 |
| *Neisseria sp.* | 0.20 | 0.96 | 0.42 | -0.16 | 0.34 | 0.16 | 0.16 | -0.14 | 0.23 | 0.00 | 0.06 | 0.36 | 0.14 | 0.13 | 0.07 | -0.04 | 0.55 | 0.15 | 1.00 | 0.01 | 0.97 | 0.27 | -0.02 | 0.02 | -0.11 | 0.02 | -0.04 | -0.04 |
| *Rhodococcus sp.* | 0.28 | 0.15 | 0.01 | 0.50 | 0.57 | 0.08 | 0.77 | 0.64 | -0.32 | 0.67 | 0.76 | 0.49 | 0.66 | -0.07 | 0.22 | 0.56 | 0.47 | 0.21 | 0.01 | 1.00 | -0.03 | 0.23 | 0.34 | 0.49 | 0.23 | 0.93 | -0.22 | 0.90 |
| *Salmonella enterica* | 0.20 | 0.93 | 0.42 | -0.12 | 0.37 | 0.15 | 0.14 | -0.22 | 0.28 | 0.08 | 0.05 | 0.35 | 0.13 | 0.19 | -0.08 | -0.10 | 0.58 | 0.26 | 0.97 | -0.03 | 1.00 | 0.30 | 0.00 | -0.01 | -0.19 | -0.01 | 0.05 | -0.09 |
| *Staphylococcus aureus* | 0.41 | 0.26 | 0.06 | 0.07 | 0.46 | 0.16 | 0.32 | -0.06 | -0.16 | 0.31 | 0.27 | 0.26 | 0.27 | -0.41 | -0.10 | 0.46 | 0.32 | 0.30 | 0.27 | 0.23 | 0.30 | 1.00 | 0.40 | 0.34 | -0.22 | 0.29 | -0.10 | 0.11 |
| *Staphylococcus capitis* | 0.27 | -0.09 | -0.27 | 0.20 | 0.37 | 0.56 | 0.33 | 0.46 | -0.07 | 0.41 | 0.33 | 0.25 | 0.28 | -0.07 | -0.10 | 0.07 | 0.13 | 0.16 | -0.02 | 0.34 | 0.00 | 0.40 | 1.00 | 0.33 | 0.14 | 0.25 | -0.36 | 0.10 |
| *Staphylococcus epidermidis* | 0.66 | 0.15 | -0.28 | 0.59 | 0.59 | 0.38 | 0.58 | 0.32 | -0.20 | 0.31 | 0.49 | 0.65 | 0.74 | -0.08 | 0.28 | 0.49 | 0.21 | -0.07 | 0.02 | 0.49 | -0.01 | 0.34 | 0.33 | 1.00 | 0.31 | 0.40 | -0.64 | 0.43 |
| *Staphylococcus lugdunensis* | 0.05 | -0.02 | 0.01 | 0.32 | 0.15 | 0.34 | 0.24 | 0.35 | -0.01 | 0.28 | 0.35 | 0.41 | 0.38 | -0.03 | 0.50 | 0.28 | -0.11 | -0.05 | -0.11 | 0.23 | -0.19 | -0.22 | 0.14 | 0.31 | 1.00 | 0.27 | -0.34 | 0.26 |
| *Streptococcus anginosus* | 0.19 | 0.17 | 0.00 | 0.52 | 0.56 | 0.05 | 0.73 | 0.59 | -0.31 | 0.65 | 0.78 | 0.50 | 0.62 | -0.17 | 0.20 | 0.55 | 0.44 | 0.25 | 0.02 | 0.93 | -0.01 | 0.29 | 0.25 | 0.40 | 0.27 | 1.00 | -0.16 | 0.85 |
| *Streptococcus pneumoniae* | -0.54 | -0.03 | 0.36 | -0.28 | -0.30 | -0.38 | -0.24 | -0.37 | 0.28 | -0.01 | -0.23 | -0.49 | -0.54 | 0.15 | -0.41 | -0.18 | 0.25 | 0.39 | -0.04 | -0.22 | 0.05 | -0.10 | -0.36 | -0.64 | -0.34 | -0.16 | 1.00 | -0.19 |
| *Streptomyces sp.* | 0.25 | 0.14 | 0.13 | 0.40 | 0.46 | -0.03 | 0.64 | 0.51 | -0.30 | 0.65 | 0.67 | 0.46 | 0.59 | -0.01 | 0.30 | 0.50 | 0.42 | 0.31 | -0.04 | 0.90 | -0.09 | 0.11 | 0.10 | 0.43 | 0.26 | 0.85 | -0.19 | 1.00 |

**Suppl. Table. 3.** Bacterial strains used in this study

| Bacterial strain | Description | Source or reference |
|---|---|---|
| *Corynebacterium accolens* | | |
| DSM 44279 | origin unknown | DSMZ |
| 63VAs_B8 | human nasal isolate | [44] |
| 83VAs_B5 | human nasal isolate | [44] |
| KPL 1824 | human nasal isolate | [45] |
| KPL 1855 | human nasal isolate | [45] |
| KPL 1996 | human nasal isolate | [45] |
| *Corynebacterium aurimucosum* | | |
| 10VPs_Sm8 | human nasal isolate | [44] |
| 48/2 | lesional site isolate | AG Peschel strain collection |
| *Corynebacterium kroppenstedtii* | | |
| 82VAs_B6 | human nasal isolate | [44] |
| *Corynebacterium propinquum* | | |
| 8VAs_B3 | human nasal isolate | [44] |
| 63VAs_B4 | human nasal isolate | [44] |
| P7-31 | human nasal isolate | this study |
| 56/2 | human nasal isolate | AG Peschel strain collection |
| 57/2 | human nasal isolate | AG Peschel strain collection |
| *Corynebacterium pseudodiphtheriticum* | | |
| 87VAs_B4 | human nasal isolate | [44] |
| 90VAs_B3 | human nasal isolate | [44] |
| P1-29 | human nasal isolate | this study |
| P2-34 | human nasal isolate | this study |
| P6-3 | human nasal isolate | this study |
| *Corynebacterium simulans* | | |
| 81MNs_B1 | human nasal isolate | [44] |
| 50MNs_Sm2 | human nasal isolate | [44] |
| 50VAs_B5 | human nasal isolate | [44] |
| 79/2 | groin isolate | AG Peschel strain collection |
| 73/2 | groin isolate | AG Peschel strain collection |
| 77/2 | groin isolate | AG Peschel strain collection |
| Corynebacterium striatum | | |
| 24/2 | human nasal isolate | AG Peschel strain collection |
| 46/2 | lesion site isolate | AG Peschel strain collection |
| 38/2 | groin isolate | AG Peschel strain collection |
| *Corynebacterium tuberculostearicum* | | |
| 87VAs_B5 | human nasal isolate | [44] |
| 12VAs_B4 | human nasal isolate | [44] |
| *Cutibacterium acnes* | | |
| 44VAs_Sa4 | human nasal isolate | [44] |

| | | |
|---|---|---|
| 50VAs_Sa1 | human nasal isolate | [44] |
| 83VAs_Sa3 | human nasal isolate | [44] |
| 87VAs_SaT9 | human nasal isolate | [44] |
| *Cutibacterium avidum* | | |
| Clone#11 | human nasal isolate | AG Peschel strain collection |
| *Dolosigranulum pigrum* | | |
| 9VAs_B4 | human nasal isolate | [44] |
| *Escherichia coli* | | |
| DH5α | K-12 derivative | New England BioLabs |
| DC10B | Δ*dcm* in the DH10B background | [27] |
| CMFI_AI 1 | axilla isolate | AG Peschel strain collection |
| CMFI_AI 2 | groin isolate | AG Peschel strain collection |
| CMFI_AI 3 | axilla isolate | AG Peschel strain collection |
| *Finegoldia magna* | | |
| 63 Vas_Sa4 | human nasal isolate | [44] |
| 83 Vas_Sa6 | human nasal isolate | [44] |
| *Klebsiella pneumoniae* | | |
| ATCC700603 | human urine isolate | ATCC |
| Lawsonella clevelandensis | | |
| DSM 45743 | human nasal isolate | DSMZ |
| *Moraxella catarrhalis* | | |
| 44VAs_Sm4 | human nasal isolate | [44] |
| 80VAs_B4 | human nasal isolate | [44] |
| *Staphylococcus aureus* | | |
| USA300 LAC | CA-MRSA isolate | [46] |
| IVK41 | human nasal isolate | [12] |
| IVK55 | human nasal isolate | [12] |
| IVK56 | human nasal isolate | [12] |
| IVK58 | human nasal isolate | [12] |
| *Staphylococcus capitis* | | |
| 10VAs_KB2 | human nasal isolate | [44] |
| 44UNs_B2 | human nasal isolate | [44] |
| 50VAs_KB6 | human nasal isolate | [44] |
| *Staphylococcus caprae* | | |
| VA18305/14 | human clinical isolate | Eppendorfklinikum, Hamburg |
| BK3880/14 | human clinical isolate | Eppendorfklinikum, Hamburg |
| *Staphylococcus epidermidis* | | |
| B1-1 | human nasal isolate | this study |
| B1-5 | human nasal isolate | this study |
| B1-10 | human nasal isolate | this study |
| B5-7 | human nasal isolate | this study |
| B5-7 Δ*sfaDAB* | deletion of *sfa* gene locus | this study |
| B5-26 | human nasal isolate | this study |
| S1-2 | human nasal isolate | this study |

| | | |
|---|---|---|
| S1-15 | human nasal isolate | this study |
| S1-27 | human nasal isolate | this study |
| S2-2 | human nasal isolate | this study |
| M14-19 | human nasal isolate | this study |
| M15-6 | human nasal isolate | this study |
| M16-32 | human nasal isolate | this study |
| W1-7 | human nasal isolate | this study |
| W1-7Δ*sfaDAB* | deletion of *sfa* gene locus | this study |
| W3-6 | human nasal isolate | this study |
| W5-7 | human nasal isolate | this study |
| W8-6 | human nasal isolate | this study |
| *Staphylococcus hominis* | | |
| 50MNs_Sa6 | human nasal isolate | [44] |
| 89VPs_B7 | human nasal isolate | [44] |
| 9VPs_KB1 | human nasal isolate | [44] |
| *Staphylococcus lugdunensis* | | |
| HKU09-01 | human skin infection isolate | [47] |
| HKU09-01 Δ*fhu* | deletion of fhu gene locus | this study |
| HKU09-01 Δ*fhu* pRB474_fhu | complementation of *fhu* gene locus; *fhu* genes encoded on pRB474 | this study |
| IVK28 | human nasal isolate | [12] |
| IVK28 Δ*fhu* | deletion of *fhu* gene locus | this study |
| *Staphylococcus pettenkoferi* | | |
| 210-70229632 | human clinical isolate | AG Peschel strain collection |
| *Mammaliicoccus sciuri* | | |
| DMS 16827 | | DSMZ |
| DMS 15613 | | DSMZ |
| 9VPs_Sm2 | human nasal isolate | [44] |
| *Staphylococcus warneri* | | |
| VA16066/14 | human clinical isolate | Eppendorfklinikum, Hamburg |
| BK6091/14 | human clinical isolate | Eppendorfklinikum, Hamburg |
| *Streptococcus pneumoniae* | | |
| ATCC 6301 | | ATCC |
| ATCC 49619 | | ATCC |

**Suppl. Table 4.** Plasmids and oligonucleotides used in this study

| Plasmid | Description | Reference |
|---|---|---|
| pIMAY | thermosensitive vector for allelic exchange | [27] |
| pRB474 | *E. coli/S. aureus* shuttle vector; constitutive active expression vector | [48] |

| Oligonucleotides | Sequence (5'-3') | Purpose |
|---|---|---|
| (1) Fhu up for | TATAGGGCGAATTGGAGCTCTTTAGTATTT TTACCTTCTGCC | Generating upstream recombinant region for *fhu* deletion |
| (2) Fhu up rev | TAGAAAGTTGGGGAATTATGTAATAAAATT CGTGTTGGAA | generating upstream recombinant region for *fhu* deletion |
| (3) Fhu dw for | CATAATTCCCCAACTTTCTA | generating downstream recombinant region for *fhu* deletion |
| (4) Fhu dw rev | GGGAACAAAAGCTGGGTACCATTATTAAGC GTATTGATGATC | generating downstream recombinant region for *fhu* deletion |
| (5) fhu chrom. Up for | CTTTCACAGATGGACTCATT | Screening for chromosomal *fhu* deletion |
| (6) fhu chrom. Dw Rev | TATTAATCTCTGTATTTTTAGG | Screening for chromosomal *fhu* deletion |
| (7) fhu comp for | TTGCATGCCTGCAGGTCGACAAATAGAAAG TTGGGGAATT | generating *fhu* genes for complementation with pRB474 |
| (8) fhu comp rev | GTACCCGGGGATCCTCTAGATTAAATAGAT TTAGCTTTATACAT | generating *fhu* genes for complementation with pRB474 |
| (9) sfaDAB ko up for | CGAGGTCGACAATAGAGAGGGATAACTAAG | Generating upstream recombinant region for *sfaDAB* deletion |
| (10) sfaDAB ko up rev | TTCTAAATGATGGTCTATCATTTAGAACGT | generating upstream recombinant region for *sfaDAB* deletion |
| (11) sfaDAB ko dw for | TGATAGACCATCATTTAGAACAATGGATTT | generating downstream recombinant region for *sfaDAB* deletion |
| (12) sfaDAB ko dw rev | ATTGGAGCTCTTATAATTAGGTCCAAGATT | generating downstream recombinant region for *sfaDAB* deletion |
| (13) Chrom sfaDAB up for | TTATAAATCATTCTTTCTATGAA | Screening for chromosomal *sfaDAB* deletion |

| | | |
|---|---|---|
| (14) Chrom sfaDAB dw rev | TTACTGGTGTGAAATGAATT | Screening for chromosomal *sfaDAB* deletion |
| (15) 473 Eco | CCTCAAGCTAGAGAGTCATTACCCC | Screening for pRB474_*fhu* positive transformants |
| (16) 473 Hind | CTGGATTTGTTCAGAACGCTCGG | Screening for pRB474_*fhu* positive transformants |
