## Supplemental Figures S1-S6 for "Siderophore piracy enables the nasal commensal *Staphylococcus lugdunensis* to antagonize the pathogen *Staphylococcus aureus*"

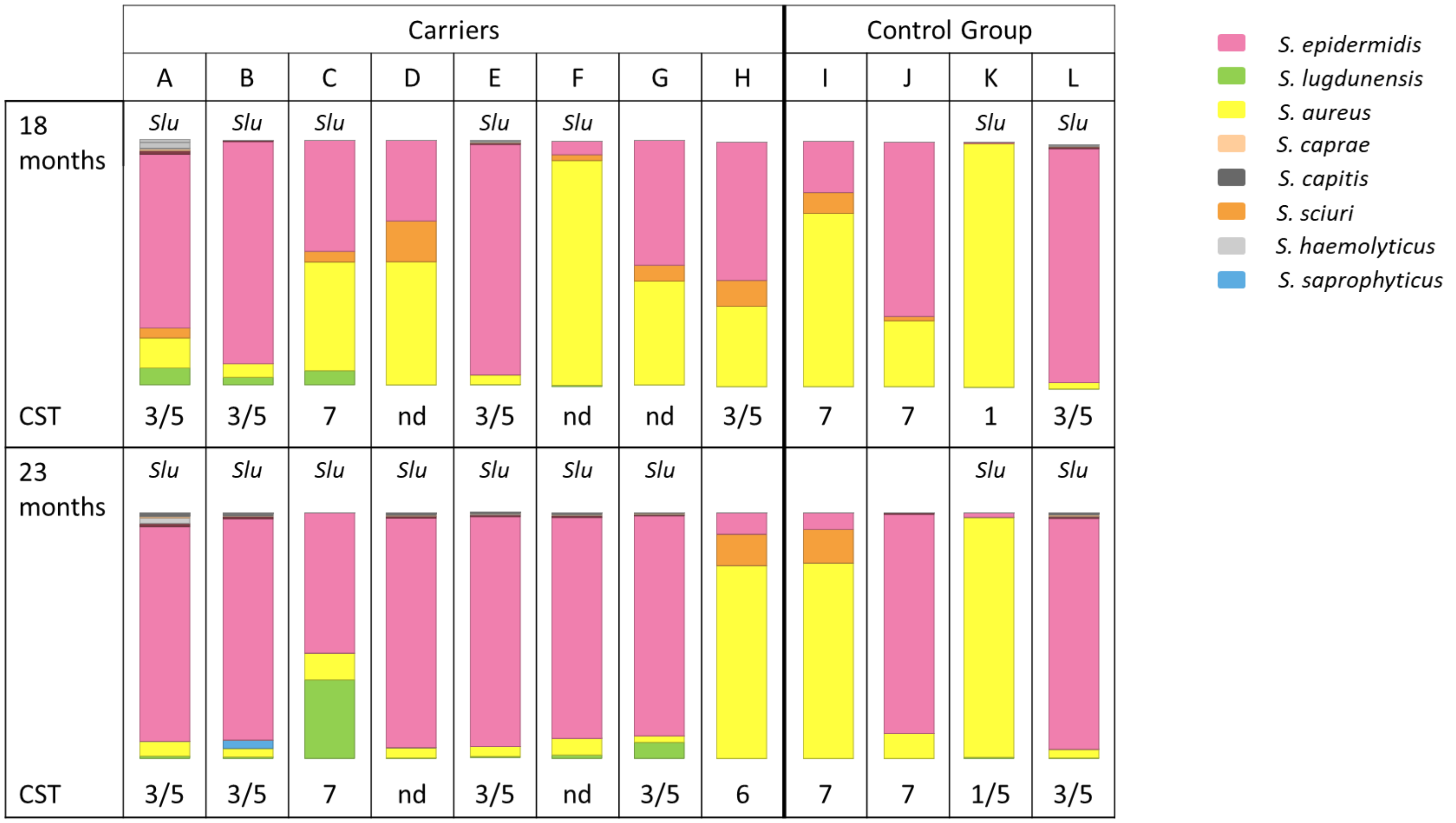


**Suppl. Fig. 1.** Microbiome profiles within the genus *Staphylococcus* at the species level. Relative proportions of staphylococcal species present in the nasal microbiomes at time points 18 months and 23 months are presented by stacked bar charts. The different bacterial genera are indicated by colours as shown in the legend on the right. The presence of *S. lugdunensis* reads in the metagenomes is indicated (*Slu*). The microbiome profiles that could be assigned to the CSTs published by *Liu et al* [11] are labelled accordingly. Profiles not fitting to one of these types are designated "nd".

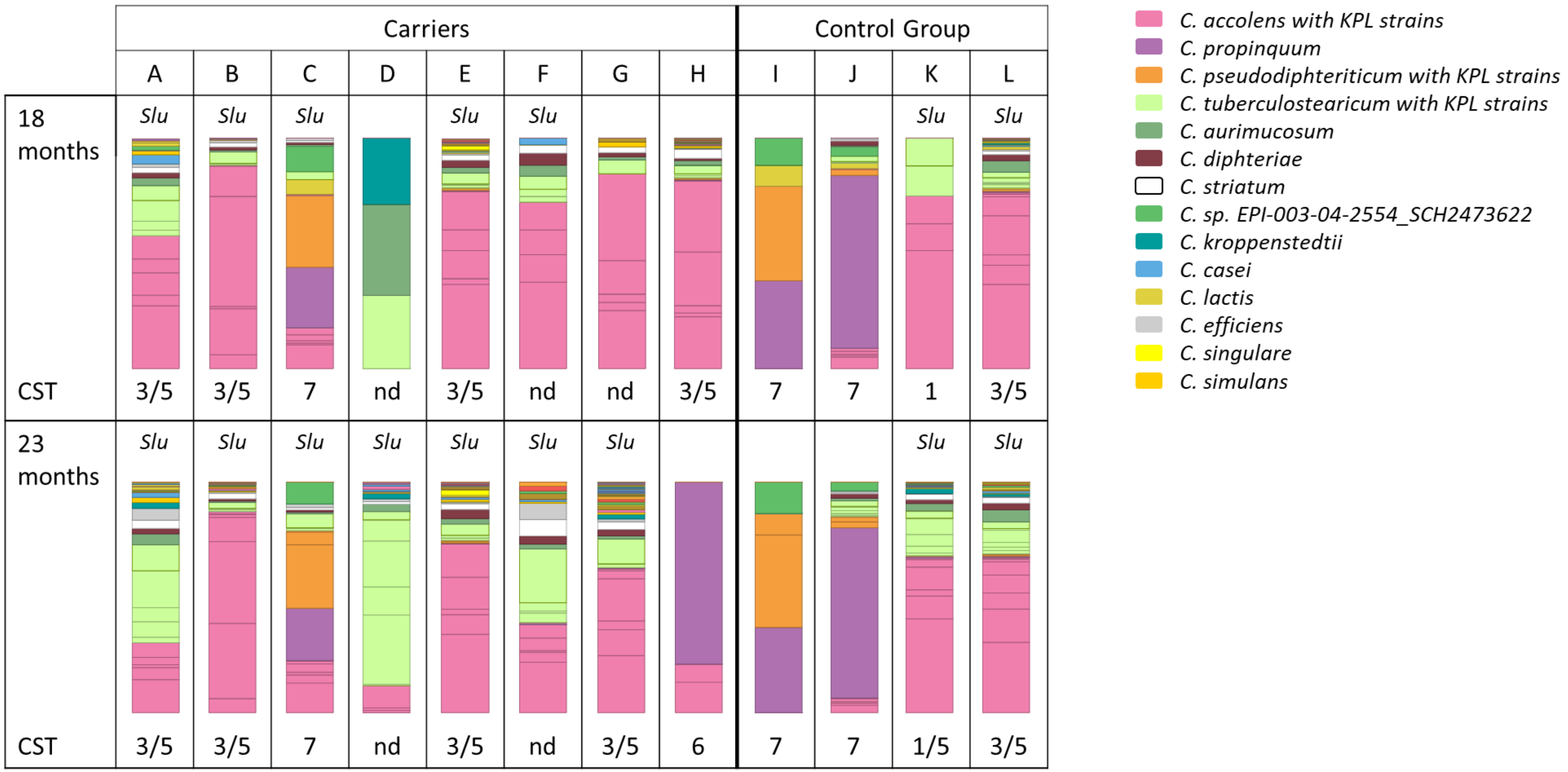


**Suppl. Fig. 2.** Microbiome profiles within the genus *Corynebacterium* at the species level. *Corynebacterium* species profiles determined in metagenome analyses at time points 18 and 23 months. Relative proportions of *Corynebacterium* species present in the nasal microbiomes are presented by stacked bar charts. The different *Corynebacterium* species are indicated by colours as listed in the legend on the right. For *C. accolens*, *C. pseudodiphteriticum,* and *C. tuberculostearicum*, different strains ("KPL strains") were determined but labeled with the same colours according to a strain assignment by the Lemon lab [45]. The presence of *S. lugdunensis* reads in the metagenomes is indicated (*Slu*). The microbiome profiles that could be assigned to the CSTs published by [11] are labelled accordingly. Profiles not fitting to one of these types are designated "nd".

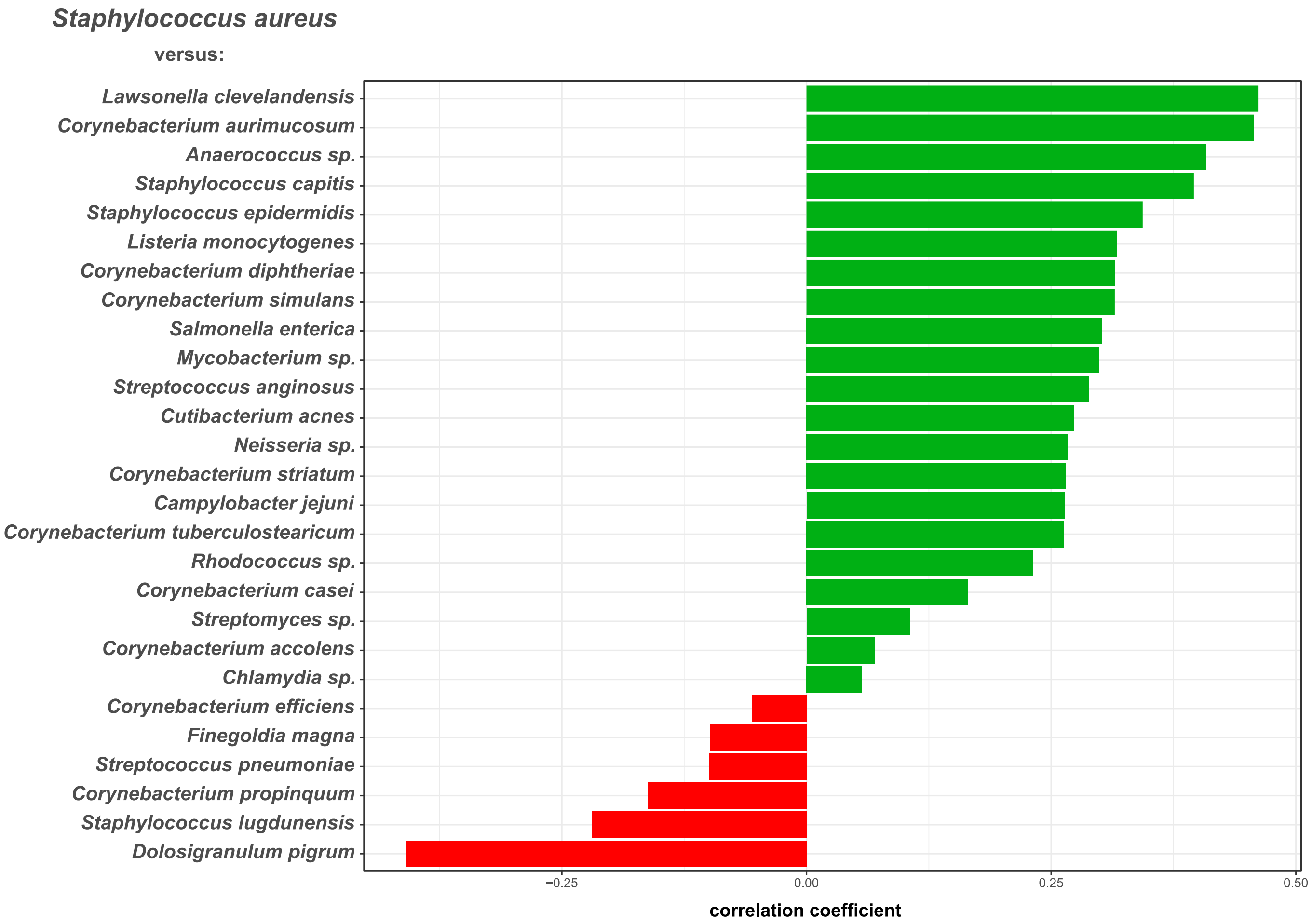


**Suppl. Fig. 3.** Correlation of *S. aureus* with other microbiome species. Correlation coefficients were inferred based on a centered log-ratio-transformed abundance table of species reads by correlation network analyses using the Namap/Pearson algorithm of MetagenoNets [20] with thresholds for prevalence and occurrence of 0.1% and 20%, respectively. Based on these parameters, we obtained correlation measures for 28 species throughout all 24 samples. Shown are the associations of *S. aureus* with other microbiome species (shown at the x-axis). Positive associations are indicated by green bars, negative associations by red bars. The corresponding Pearson correlation coefficients are indicated at the x-axis.

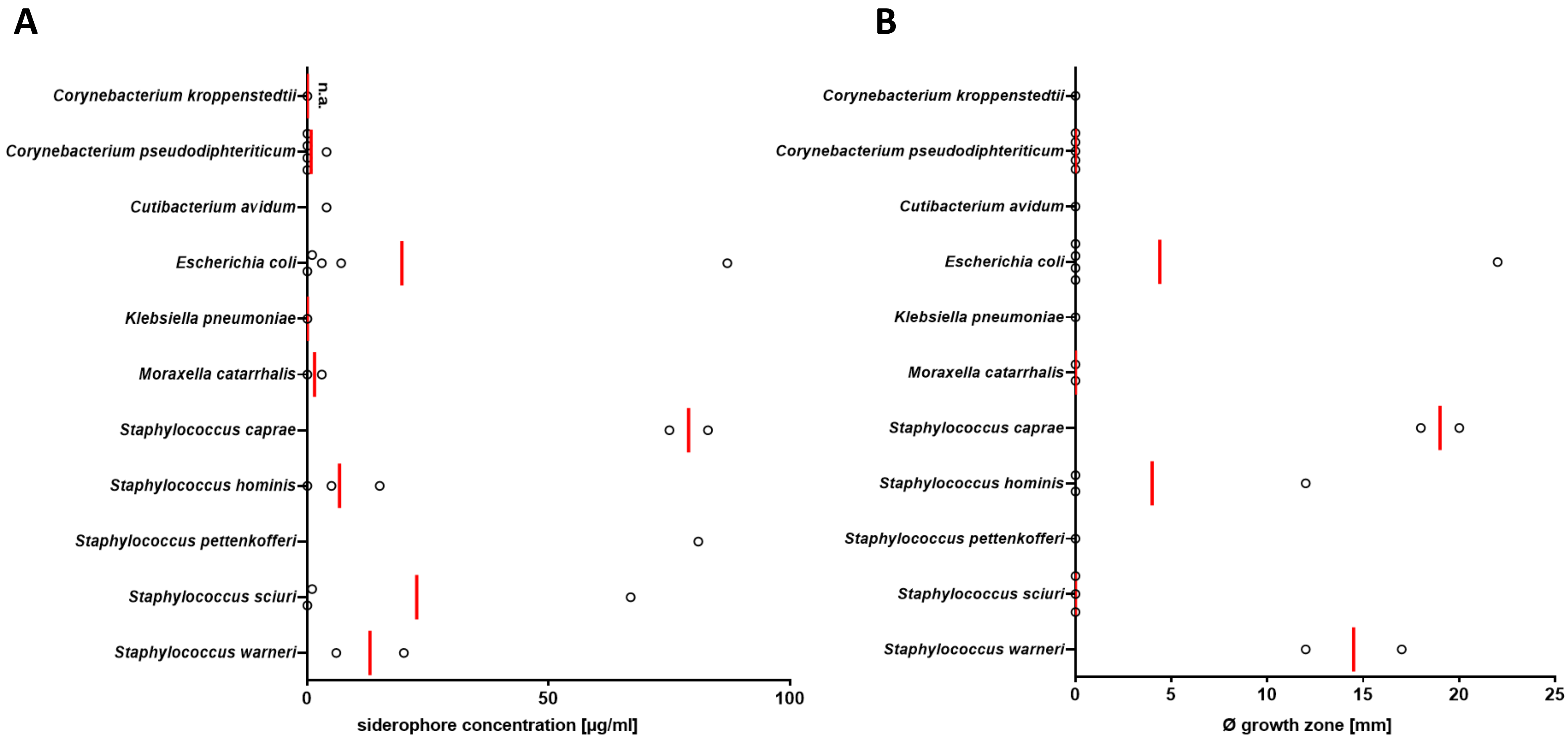


**Suppl. Fig. 4.** Nasal bacterial species producing siderophores have the capacity to support growth of *S. lugdunensis*. (A) Siderophore concentration in spent media from nasal bacteria after three days of incubation detected via SideroTec Assay$^{TM}$ kit. Siderophores could be detected in spent media from staphylococcal species and *E. coli*; siderophore detection was not assessable for *C. kroppenstedtii* due to the essential presence of Tween 80 in growth medium that interacts with the reaction solutions provided by the SideroTec Assay$^{TM}$ kit. (B) Occurrence and sizes (diameter in mm) of growth zones of *S. lugdunensis* promoted by siderophore-containing spent media from nasal bacteria. Staphylococcal species and *E. coli* support *S. lugdunensis* growth under iron-restricted conditions. Data points represent average values of three independent biological replicates, and vertical red lines show means of each species.

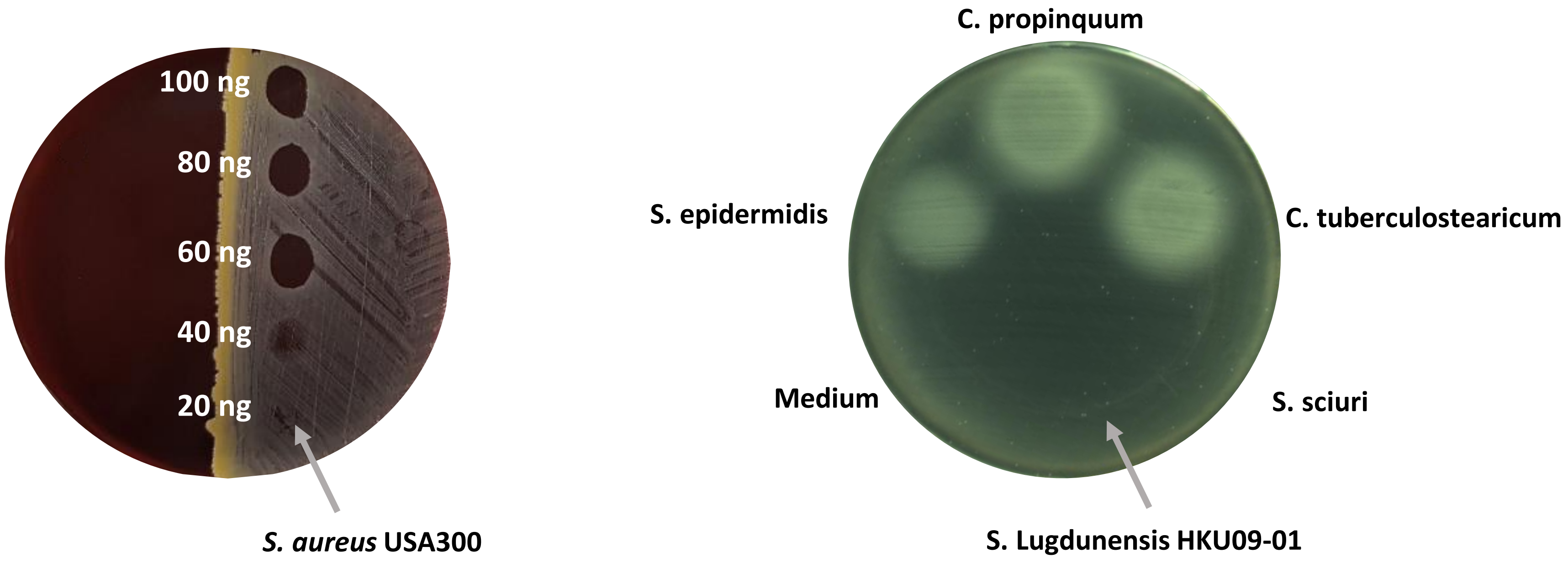


**Suppl. Fig. 5.** Representative pictures of lugdunin susceptibility determination of nasal bacteria and *S. lugdunensis* growth promotion by siderophore-containing spent media from nasal bacteria. (A) Lugdunin susceptibility determination by the example of *S. aureus* USA300. 2 µl containing different amounts of lugdunin (0 to 100 ng) were spotted onto lawns of bacteria on sheep blood agar plates, resulting in zones of inhibition for lugdunin-susceptible bacteria. The spot with the lowest amount of lugdunin showing a clear inhibition was used to determine lugdunin susceptibility. (B) Growth promotion of *S. lugdunensis* HKU09-01 by siderophore-containing spent media of nasal bacteria. Iron-restricted BHI agar supplemented with 10% horse serum (as transferrin source) were inoculated with a lawn of *S. lugdunensis*. Spent media of different bacteria were spotted onto *S. lugdunensis* lawns resulting in zones of enhanced growth when siderophores were present. Diameters were measured and noted.

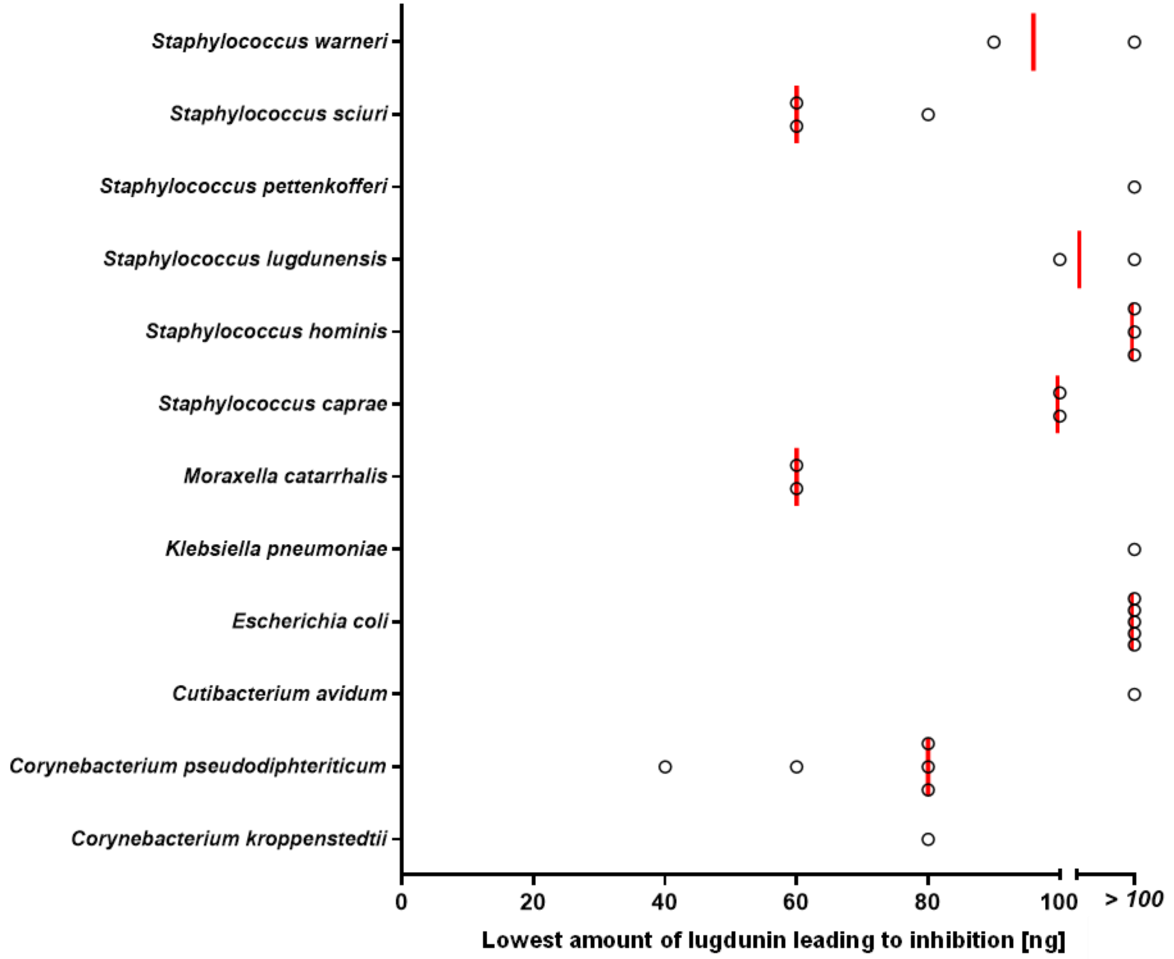


**Suppl. Fig. 6.** Lugdunin susceptibility of nasal bacterial species. Different amounts of lugdunin (0 to 100 ng in 2 µl) were spotted onto lawns of nasal bacteria, resulting in zones of inhibition. Data points represent average values of three independent experiments, and vertical red lines show medians of each group.
